# The SorCS2-derived macrocycle TT-P34 drives neuroprotection in animal models of neurodegeneration

**DOI:** 10.1101/2025.09.17.676723

**Authors:** Anders Dalby, Mathias Kaas, Johan Palmfeldt, Mads Graversgaard, Sanne Nordestgaard, Christian R. O. Bartling, Neil Benson, Emily Roashan, Søren L. Pedersen, Larry C. Park, Simon Glerup, Kristian Strømgaard, Keld Fosgerau, Simon Mølgaard

**Author notes:** Corresponding authors; mail.

## Abstract

Mitochondria are critical for sustaining the high energy demands of neuronal activity and their dysregulation is a hallmark of neurodegeneration. Targeting pathways of neurotrophic signaling is a well-established therapeutic strategy to enhance mitochondrial function and mitigate neurodegeneration. The VPS10p domain receptor, SorCS2, has recently emerged as a receptor with neurotrophic signaling capabilities. Here, we design and develop novel SorCS2-derived macrocyclic peptides mimicking receptor activation *in vivo*. We show that SorCS2-peptides enhance both neurotrophic support and boost metabolism by activating CREB and AMPK in a CAMKK2-dependent manner. This leads to upregulation of the key transcription factors PGC1α and TFEB and consequentially mitochondrial biogenesis. Furthermore, we show that the lipidated SorCS2 macrocycle, TT-P34, rescues motor behavioral deficits and preserves synaptic and mitochondrial signatures in the zQ175 mouse model of Huntington’s Disease. In addition, treating a MPTP-induced mouse model of Parkinson’s Disease leads to amelioration of behavioral deficits and reduction of dopaminergic loss. Finally, we demonstrate that TT-P34 crosses the blood-brain barrier in non-human primates, and estimate human therapeutic dosing by pharmacodynamic modelling. Together, our findings support the use of TT-P34 as a novel disease-modifying therapy targeting SorCS2-receptor signaling to prevent neurodegeneration.

## INTRODUCTION

Neurodegenerative diseases (NDDs), including Alzheimer’s (AD), Parkinson’s (PD), Huntington’s Disease (HD), and amyotrophic lateral sclerosis (ALS), are chronic conditions of the brain affecting millions of people worldwide. As the aging population continues to grow, these figures are expected to increase, with projected dementia cases rising from 21 million in 2025 to 37 million by 2050 [1]. Despite extensive efforts, available therapies are focused on symptomatic relief.. However, many novel drug candidates are in clinical development addressing new mechanistic targets inspiring optimism for future treatments.

Neurotrophins, including nerve-growth factor (NGF), brain-derived neurotrophic factor (BDNF) and others, have long been considered a promising treatment approach in AD, PD, ALS and HD due to their regenerative and neuroprotective features. Patients suffering from NDDs exhibit lower BDNF levels in both serum and brain [2–9] and analysis of post-mortem tissues have linked cyclic AMP response element-binding protein (CREB), the central transcription factor targeted by NGF and BDNF-signaling, as a critical transcription factor affected across NDDs [10–16]. Unfortunately, clinical trials using exogenous administration of recombinant neurotrophins have failed primarily due to limited blood-brain barrier penetration and their poor diffusion across the brain [17–19]. This has prompted a shift towards gene delivery methods and, while a phase 1 clinical trial of AAV2-BDNF gene therapy for early AD is currently ongoing, the overall lack of success has sparked an interest in targeting the BDNF/TrkB pathway through alternative strategies including the use of monoclonal antibodies, small molecules, and peptides [20–23].

The Sortilin Related VPS10 Domain Containing Receptors 1-3, SorCS1-3, are well established co-receptors of neurotrophin receptors TrkB and p75NTR (in addition to their respective neurotrophins, pro and mature NGF and BDNF) important for their synaptic targeting, internalization and downstream signaling [24–28]. These receptors have additionally been genetically and functionally implicated in NDDs including AD, HD and ALS [29–35].

We previously identified a triple-serine signaling motif within the intracellular domain of SorCS2 capable of transducing downstream signaling independent of BDNF[36]. Utilizing this knowledge, we generated cell-penetrating peptides capable of activating CREB leading to transcriptional activation of mitochondrial genes of oxidative phosphorylation of which dysfunction is a common pathological hallmark in NDDs.

Here we outline the development of two novel CREB-activating SorCS2-derived macrocyclic peptides with different pharmacokinetic profiles. We demonstrate that once-daily subcutaneous (s.c.) administration in rodents ameliorates behavioral deficits and protects against synaptic dysfunction in two models of neurodegenerative diseases (HD and PD) by engagement with mitochondrial and lysosomal systems demonstrated through proteomic and metabolomic analysis. Together, this outlines the promising therapeutic potential of targeting the SorCS2/BDNF/TrkB pathway in neurodegenerative diseases.

## MATERIALS AND METHODS

### Animal handling and husbandry

C57BL/6j BomTac wild type (WT) mice were purchased from Taconic (pharmacodynamics studies) or GemPharmatech Co, LTD (MPTP study). Experiments for pharmacodynamics were carried out at Aarhus University and approved by the Danish Animal Experiments Inspectorate under the Ministry of Justice (Permit 2011/561-119). The execution of the experiments adhered to both institutional and national regulations. Cages were cleaned every week and supplied with bedding and nesting material, a wooden stick, and a metal tunnel. For the MPTP study, the animals were housed in a temperature (22 ± 3 °C) and humidity (50 ± 20%) controlled AAALAC accredited facility at WuXi AppTec, Pudong, Shanghai. In all studies the mice were provided with food and water ad libitum and kept on a 12-hour light/dark (6:00 am /6:00 pm) cycle.

The zQ175DN mice (zQ175DN KI, B6J.zQ175DN KI with ∼CAG 190) and 20 aged match wildtype (WT) littermates were received from CHDI Foundation, Inc. All zQ175 animal experiments were carried out at Naason Science, Inc.,, Cheongju-si in accordance with the Ministry of Food and Drug (MFDS) guidelines for the care and use of laboratory animals and approved by Naason Science in accordance with all applicable IACUC (IACUC number: KBIO-IACUC-2021-157-1) and AAALAC guidelines.

Animal studies in rats (Sprague Dawley) were performed at Aptuit, Verona under Italian Legislative Decree no. 26/2014 (authorization issued by the Italian Ministry of Health) or Biotest Facility, Aarhus under license 2021-15-0201-01064 with the Animal Experiments Inspectorate supplied by Janvier Labs. Bedding was changed once a week, air exchanged approximately 12 times per hour in the stable and temperature was controlled via the ambient ventilation system to be between 20-24 °C. The rats were provided with diet and water ad libitum and light cycle was 12-hour dark and 12-hour light (lights on 6.00).

Male Beagle Dogs supplied by Marshall Bioresources, were used for pharmacokinetics of TT-P34. Experiments were carried out at Aptuit, Verona under European Directive no 2010/63/UE. All studies was conducted in accordance with national legislation. Dogs were provided with diet and water ad libitum, had access to toys with certificates of analysis, held in stable with 19-21 °C temperature with fluorescent lighting from approximately 06:00 to 18:00 hours daily. The dogs was monitored 24 hours a day.

The non-human primate (Macaca fascicularis (PNH)) study was performed at Motac Neuroscience, Bordeaux, and in agreement with the European Communities Council Directive (2010/63/EU) on the protection of animals used for scientific purposes. The study was approved by the local Institutional Animal Care and Use Committee (IACUC, Comité d’Ethique en Expérimentation Animale de Bordeaux - CEEA50) with a generic but not study-specific registered under number #44183. All NHPs had free access to pellets specific for old world monkeys (SDS) with banana flavor, with fresh fruit and vegetables distributed daily. Drinking water was available ad libitum. Toys and mirror were provided in the home cages. The radio played in the husbandry area for one hour once a day.

### Culturing

For culturing and preparation of murine primary neuronal cultures see Dalby et al. 2025. The human iPSC-derived glutamatergic and GABAergic neurons were acquired from Bit.Bio and cultured and maintained according to their protocols. All assays on human iPSC-derived neurons were carried out at 14 days in vitro (DIV). Huntington’s patient derived fibroblasts (GM04476) were acquired from Coriell Cell Repositories and maintained in DMEM (ATCC, ATCC-30-2002) containing 15% FBS (Sigma, F9665) and Pen/Strep (Gibco, 15070063) at 37 °C.

### STO-609 assay

Murine primary cortical neurons were seeded in precoated (Poly-L: Sigma #P1524 and laminin: Invitrogen, 23017-015) 12-well plates at a density of 5×10^5^ cells/well. At 7DIV half of the neurons were treated with STO-609 (CaMKK2 selective inhibitor dissolved in DMSO, Selleckchem, S8274) at a final concentration of 5uM and half received DMSO only. After 1 hour of incubation at 37 °C, the media was removed and the neurons were treated with prewarmed Neurobasal-A Medium (Gibco, #10888022) containing either TT241 or TT241c and STO609 or no additives and incubated for 10 min. at 37 °C. After incubation, the cells were lysed on ice in KINEXUS special lyse-buffer (20 mM MOPS, 2 mM EGTA, 5 mM EDTA, 50 mM Sodium fluoride, 60 mM b-glycerophosphate (pH 7.2), 25 mM sodium pyrophosphate, 2.5 mM sodium orthovanadate, 50 nM phenylarsine oxide, 1% triton X-100 and 0.05% sodium dodecylsulphate (SDS)) containing Pierce Protease Inhibitor Mini Tablet (ThermoFisher, A32953) and DTT and subsequently sonicated in a cup horn at 4 °C for 1 min. Lysates were collected and stored at −20 °C until further processing (see Protein analysis).

### Mutated HTT assessment

One day prior to the experiment, HD patient derived fibroblasts were seeded in a 96-well at a density of 3-5×10^4^ cells/well in 100 µL DMEM containing 15%FBS and P/S at at 37 °C. At the day of the experiment the cells were stimulated to a final concentration of 1 µM of TT241 or 1 µM of TT-P34 once by direct addition of 10 µl of peptide stock solution in DMEM (no additives). 24 hours after stimulation, the cells were lysed on ice in RIPA buffer (50 mM Tris pH 7.4, 150 mM NaCl, 1% Triton X-100, 2 mM EDTA, 0.5% Sodium-dexoycholate, 0.1% SDS) containing Pierce Protease Imhibitor Mini Tablet (ThermoFisher, A32953) and left for shaking for 30 min at 300 rpm. Lysates were collected and stored at −20 °C until further processing (see Protein analysis).

### Protein analysis

For SDS-PAGE and Western blotting see Dalby et al. 2025. Primary antibodies used for western blotting were 1:1000 rabbit anti-human/mouse phospho-CREB S133 (Cell Signaling Technology #9198), 1:1000 rabbit pAb phospho-AMPK T172 (Cell Signaling Technology #2531S), 1:1000 rabbit pAb anti-TFEB (Thermo Fisher Scientific, A303-673A), 1:1000 rabbit pAb anti-PGC1α (Novus Biologicals, NBP1-04676), 1:1000 rabbit mAb anti-BDNF (Abcam, ab108319), 1:1000 mouse mAb anti-total HTT antibody (MAB2166, Sigma), 1:1000 mouse mAb anti-polyQ specific (MABN2427, Sigma), 1:5000 mouse anti-mouse beta-actin (Sigma #A5441). Secondary antibodies used for western blotting were anti-rabbit IgG, HRP-linked Antibody (Cell Signaling Technology, 7074S), and Rabbit Anti-Mouse (Agilent, P026002-2).

Simple Western was run on Abby (Bio-Techne, 004-680) following their protocols on a 12-230 kDA Separation Module (#SM-W004, Bio-Techne). Lysates were spun at 13000 rpm for 10 min at 4 °C. Supernatant from lysates were combine 5x Fluorecent Master Mix with parts lysate (4:1), vortexed and heated for 5 min at 95 °C with 300 rpm shaking. Plates were loaded in addition to primary and secondary antibodies, wash buffer, luminol-peroxide, biotinylated ladder and RePlex reagent. Primary antibodies used were 1:20 rabbit pAb anti-D1R (Abcam, ab20066), 1:20 rabbit mAb anti-DARPP32 (Abcam, ab40801), 1:20 rabbit pAb anti-SCN4B (Abcam, ab80539) and 1:100 mouse anti-mouse beta-actin (Sigma #A5441). Secondary antibodies used were anti-Mouse Secondary antibody (#042-205, Bio-Techne), anti-Rabbit Secondary HRP antibody (#042-206, Bio-Techne), 20x Anti-Rabbit HRP Conjugate (#043-426, Bio-Techne).

### MTT assay

5×10^4^ cells/well murine primary cortical neurons were plated per well in a 96-well plate. At 7, 9 and 11 DIV half the media was changed with new media (Neurobasal-A Medium with B-27™ Supplement (50X), serum free, ThermoFisher, 17504044) containing a dose-range of TT241. At 12DIV, half the media was removed and 10 µl MTT (3-[4,5-dimethylthiazol-2-yl]-2,5-diphenyltetrazolium bromide, Sigma, 11465007001) were added to each well to give a finale concentration of 0.5 mg/mL. To control for background absorbance 10 µl of MTT were added to well containing 100µl media only. The cells were incubated for 3-4 hours at 37 °C. After incubation the media was carefully removed and 100 μL of solvent (50% DMSO, 50% EtOH v/v) were added to each well and the plate left for shaking for 30 minutes (covered in alu-foil). Finally, absorbance was measured at 570 nm and 650nm for wavelength correction.

### In vivo pharmacodynamics and kinetics

C57BL/6j BomTac wild type (WT) mice or SD rats were allocated to room of injection and dissection minimum 24 hours prior to the experiment. Mice were injected by subcutaneous administration with doses demonstrated in figures in formulation 4.38 mM L-His, 140mM NaCl, 0.2% Tween-20 and 1500IU hyaluronidase (Sigma, 37326-33-3) and rats in PBS formulation. Each cage receives all treatments as to ensure minimal cage-cage bias. After indicated timepoints in figures, the mice were sacrificed and hippocampus and striatum dissected on ice, put into 2 mL Eppendorf tubes and snap frozen in liquid nitrogen. Tissue was stored at –80 °C awaiting further processing. All tissues were lysed in RIPA buffer (see above) containing containing Pierce Protease Inhibitor Mini Tablet using TissueLyser II (Qiagen, #85300) and Tungsten Carbide Beads, 3 mm (QIAGEN, 69997) for 2 min., 50 Hz at 4 °C. Samples were spun down for 10 min. at 4°C at 14800 rpm. Supernatant was stored at –80 °C awaiting further processing. For pharmacokinetic studies, brain tissue used for peptide measurements were externally washed with physiologic solution and collected in Precellys Vials and stored at −80 °C. All blood samples were collected into potassium EDTA tubes, centrifuged (3000g for 10 min at approximately 4 °C) and stored at −80 °C.

### zQ175 study

The study examined male and female animals, and similar findings are reported for both sexes. zQ175 mice were injected once daily by subcutaneous administration with vehicle (formulation only) or TT-P34 (0.2mg/kg) or TT241 (13mg/kg) in formulation 4.38 mM L-His, 140mM NaCl, 0.2% Tween-20 and 1500IU hyaluronidase (Sigma, 37326-33-3) from 3 months of age until 12 months of age. Age-matched wildtype (WT) littermates were injected with vehicle only. Body weight was measured every week starting at 3 months of age until the end of the study. Behavioral tests (rotarod and transverse beam tests) and HCA were performed at 3 (pre-treatment), 6, 9, and 12 months of age. MRI of the brain was taken at 9 and 12 months of age. CSF and brain tissue was collected and snap for further analysis (proteomics, metabolomics and western techniques). Plasma was also collected as previously described.

#### Rotarod

Mice were tested during the diurnal phase over 2 consecutive days at 3 (pre-treatment), 6, 9 and 12 months of age. Each daily session included a training trial of 5 min at 4 RPM on the rotarod apparatus (AccuScan Instruments, Columbus, USA). One hour later, the animals were tested for 3 consecutive accelerating trials of 6 min with the speed changing from 0 to 40 RPM over 360 seconds and an inter-trial interval at least 30 min. The latency to fall from the rod was recorded. Mice remaining on the rod for more than 360 seconds were removed and their time scored as 360 seconds.

#### Transverse Beam Test

Mice were tested during the diurnal phase 3 (pre-treatment), 6, 9 and 12 months of age. The beam test was adapted from the procedure described by Fleming et al [37]. Motor performance was measured with a novel beam test adapted from traditional beam-walking tests. Animals were trained to traverse the length of the beam starting at the widest section and ending at the narrowest, most difficult section. Animals received 2 days of training before testing, and all training was performed without the mesh grid. On the first day, animals received two assisted trials, which involved placing the animal on the beam and positioning the home cage in close proximity to the animal. On day 2 of training, animals were required to run five trials. To increase difficulty further, on the day of the test, a mesh grid (1 cm squares) of corresponding width was placed over the beam surface leaving a ∼1 cm space between the grid and the beam surface. Animals were videotaped while traversing the grid-surfaced beam for a total of five trials. Videotapes were viewed and rated in slow motion for errors, number of steps made by each animal, and time to traverse across five trials by an investigator blind to the treatment group. Error of steps and the time to traverse were measured for WT and zQ175 mice across all five trials and averaged.

#### Home Cage Analysis

zQ175 and WT mice at the age of 2.5 months were placed in home cages at N=4/cage in random groups. RFID chips were introduced to each animal for the purposes of this monitoring. The home cage monitoring was performed on a 48-hour rotation at 3 (pre-treatment), 6, 9 and 12 months of age. All groups of mice (n=8 per group, n=4 per gender) were home cage monitored with ActualHCA™ devices for 48 hours per cage. The following parameters were measured: moving time, isolated time, peripheral time, in center zones time, climbing time, drinking time, moving speed, moving distance, isolation/separation distance, peripheral distance, in center zones distance and body temperature. The parameters measurements were first processed using the HCA software followed by the algorithms developed by Naason Science using Matlab software (USA).

#### MRI scans

MRI scans were performed in all groups (n=6 per group, male 3, female 3) 12 months of age (MRI). MRI acquisitions were performed using a horizontal 4.7T MRI magnet (Bruker Biospin GmbH, Ettlingen, Germany; different apparatus was used for each timepoint). Mice were anesthetized using isoflurane, fixed to a head holder and positioned in the magnet bore in a standard orientation relative to gradient coils. Anatomical images were acquired using a TurboRARE sequence with TR/TE = 2500/36 ms, matrix size 256 × 256, FOV 20.0 × 20.0 mm2, 19 contiguous 0.7 mm thick slices and 8 averages.

#### Neuronal NfL measurements in CSF & Plasma

Concentrations of neurofilament light chain protein (Nf-L) in the CSF were determined using the Simoa Neuronal 4PlexA assay (cat# 102153, Quanterix). Paramagnetic carboxylated beads (Quanterix Corp, Boston, MA, USA) were coated with mouse Nf-L antibody and incubated for 35 min with the sample and a biotinylated mouse anti–Nf-L antibody in the Simoa instrument (Quanterix). The average number of enzymes per bead (AEB) of samples was interpolated onto the calibrator curve constructed by AEB measurements. Plasma NfL levels were determined using R-PLEX Human Neurofilament L Assay (Mesoscale Discovery, K1517XR-2).

### Proteomics

Protein pellets from striatal samples were treated by filter aided sample preparation[38]. The subsequent LC-MS based label-free proteomics was essentially performed as previously described[39]. Peptide mixtures were purified with PepClean™ C18 Spin Columns and dried on a miVac Duo Concentrator (Genevac). Samples were stored at −20°C until LC-MS/MS analyses. Peptide samples were analyzed by nano liquid chromatography (Easy-nLC 1200, Thermo Scientific, Waltham, MA)-tandem mass spectrometry (Q-Exactive HF-X Hybrid Quadrupole Orbitrap, Thermo Scientific, Waltham, MA) through data dependent acquisition. Proteins were identified and quantified using MaxQuant (version 1.5.3.30). False discovery rate threshold of protein identification was set to 0.005. Differential alteration analyses were performed for the proteins with quantitative values in at least 5 out of 9 of the replicates of each sample group and at least 24 of 36 replicates overall. Differentially altered proteins (DAP) were defined as proteins with fold change (FC) higher than 1.5 and p<0.05 (corrected for multiple comparison, i.e. the 5 t-tests).

### Metabolomics

The metabolite samples were analyzed essentially as previously described[40]. The metabolite samples were cleared from proteins by protein precipitation by 80% methanol and analyzed by LC-MS on Vanquish LC and Q Exactive Plus Orbitrap MS, both from Thermo Scientific. 10 µl of sample was injected and the LC gradient constituted of a 10 min gradient with 3-23% methanol, followed by washing and re-equilibration. Both buffer A (LC-MS grade water) and B (LC-MS grade methanol) contained 0.2% formic acid. The analytical LC column was a 100 mm long Biphenyl column (Accucore, from Thermo Scientific) with inner diameter of 2.1 mm. The MS analyses were performed in positive MS mode at an electrospray voltage of 3500 V, with scanning from 70-1050 m/z. Stepwise fragmentation was performed with normalized collision energy levels of 20, 40 and 60. The MS was operated at high resolution (70,000) and accurate mass (<5 parts per million) to assure high analytical selectivity. Compound Discoverer 3.3 (Thermo Scientific) was used for identification and quantification of the compounds. The identification nodes were 1) predicted composition and the databases 2) mzCloud (endogenous metabolites), and 3) Chemspider (Human Metabolome Database, KEGG). The maximal allowed mass deviation between experimental MS data and database values was 10 parts per million.

### MPTP study

The study examined male mice because male animals exhibited less variability in phenotype. It is unknown whether the findings are relevant for female mice. MPTP was injected at 30 mg/kg per day from 3 to 10 days, started test compound treatment at day 1 daily for up to 10 days. At day 9 and 10, 6 hours after MPTP dosing, behavioral test was performed. At day 10, after grip strength measurement, animals were sacrificed and plasma and CSF were collected and stored at −80°C. Mice were then intracardially perfused with ice-cold saline. Brain tissues were collected for TH IHC.

#### Behavior

On day 9 of the study, 6 hours after MPTP injection, the animals were placed in the open field to detect changes in motor behavior (total distance traveled and vertical counts) within 10 minutes. On day 10 of the study, 6 hours after MPTP injection, for grip force measurements, mice were allowed to grip the metal grids of a grip meter with their forelimbs, and then they were gently pulled backwards by the tail until they could no longer hold the grids. The average grip strength observed in 10 trials were recorded and calculated.

#### Immunostaining of tyrosine hydroxylase

Mouse brain tissues were processed using standard histological methods and embedded in paraffin. Coronal sections of mouse brain tissue were embedded in paraffin. Two slide sections were cut, 4um thickness, at Bregma −3.08 mm and Bregma −3.52 mm. The section were deparaffinized and dehydrated. Slides were immersed in 3% hydrogen peroxide solution at room temperature. Antigen retrieval buffer of Citrate (pH6.0) was used.. To avoid nonspecific staining, the sections were incubated in blocking serum (DAKO, X0909) for 15 mins at room temperature, followed by using primary tyrosine hydroxylase antibodies (Abcam, ab137869) in dilution 1: 900 for 1 hour. Then secondary goat polyclonal antibodies conjugated to HRP (DAKO#K4003) were added. For image analysis of neurons, TH-stained sections were used and scanned by Leica Aperio GT450 Scanner. Images were opened with HALO, TH-positive cells of SNpc on both sides were counted.

### Non-human primate stereotaxic surgery

The animals were fasted overnight prior to surgery. On the day of surgery, each animal was given atropine SO4 (Atropine Sulphate Aguettant) at 0.05 mg/kg intramuscularly (i.m.) and diazepam (Diazepam TVM 5 mg/mL, 0.5 mg/kg, i.m.). At least 10 minutes later, each animal was anaesthetised with a mixture of Ketamine HCl (Ketamine® 1000 Virbac, 10 mg/kg, i.m.) and xylazine HCl (Rompun® 2% Bayer, 15 mg/kg, i.m.). The non-steroidal anti-inflammatory drug Meloxicam (Metacam®) was then given at 0.2 mg/kg subcutaneously (s.c.). Animals were placed on a heating pad with a second pad placed on top of the animals when required as indicated by a drop in the animal’s core body temperature (measured by probed placed rectally). Each animal was positioned for surgery in a stereotactic apparatus. The surgical site was prepared for aseptic surgery by initially wiping the area with sponges soaked in 70% isopropyl alcohol scrub, which was allowed to dry, followed by application of DuraPrepTM (or similar solution), which was also allowed to dry. Topical anaesthesia was applied. The animal was appropriately draped for strict aseptic surgery. After making a small craniotomy, without damaging the dura matter, a ventriculographic cannula mounted on a glass syringe was introduced into the anterior horn of the lateral ventricle and a contrast medium (0.1 mL of Omnipaque, Nycomed, Norway) injected. A stereotactic atlas was used for precise adjustment before insertion into the skull. The precise position of the anterior commissure was determined from ventriculography. The actual position of the right putamen and the lateral ventricle was defined by combining the ventriculography-defined position of the anterior commissure and a stereotactic population-based historical atlas of the basal ganglia. The area of the craniotomy was cleaned with sterile saline. Placement of a single guide cannula into the right putamen (n=1) and the lateral ventricle (n=1) for microdialysis was performed using a dedicated CMA Kopf-compatible cannula holder. The plastic pedestal of the cannula was secured to the skull with stainless steel self-tapping slot pin screws (Stryker, France). The wound was closed in layers with a continuous pattern of absorbable suture material. The skin was closed with an appropriate size of absorbable suture material, placed in a subcuticular pattern. The animal was then allowed to recover from anaesthesia and kept warm until it has regained the righting and swallowing reflexes. The animals were under the care of experienced technicians with ready access to experienced veterinary support. All surgical incisions were observed for signs of infection and inflammation and general observations was carried out to assess well-being post operation.

### Non-human primate microdialysis

Animals were seated in a restraining chair to which they had been acclimatized. Probe (CMA 11, PES, 260 µm outer diameter, 4 mm membrane length; MWCO 55 KDa) was inserted 1 hour prior to baseline sampling. Probes were perfused at a constant flow rate (0.5 µl/min) by means of a microperfusion pump (CMA 402, Phymep, Paris, France) with artificial cerebrospinal fluid (aCSF) containing (in mM): 153.8 Cl-, 147 Na+, 2.7 K+ 0.85 Mg2+, and 1.2 Ca2+, ∼ pH 6. A polyethylene tube connected the inlet of the microdialysis probe with a microinjection pump and the outlet with a collection vial. A stable baseline, defined as three consecutive samples in which the dialysates contents varied by less than 10%, was then obtained 120 min after the beginning of the perfusion as planned. After baseline sampling, TT-P34 (2mg/kg) was administered by s.c. administration. One sample was collected every 20 min on ice, frozen in dry ice and stored at −70°C. Plasma was additionally sampled for up to 192 hours post administration. In a separate study, TT-P34 (2mg/kg) was administered intravenously and plasma was sampled for up to 192 hours post administration.

### Statistical and functional enrichment analysis

Data are presented as mean ± SEM. Statistical analysis was performed using, one-way ANOVA, two-way ANOVA followed by post hoc analysis using Student’s t-test unless mentioned otherwise. Nonparametric tests like Mann-Whitney and Kruskal-Wallis tests were used when N was too small or data did not follow Gaussian distribution. The difference was considered significant when p< 0.05.

Functional enrichment analyses were performed by MetaboAnalyst 6.0 (https://www.metaboanalyst.ca/) with pathway cluster group threshold at score 1.3 (corresponding to p<0.05).

## RESULTS

### The SorCS2-derived cyclic peptide TT241 activates mitochondrial pathways

We previously designed a SorCS2 phosphomimetic peptide, here denoted TT221, constituting a 21 amino acid (aa) SorCS2-derived peptide attached to an 11 aa Tat-like cell-penetrating moiety (to allow intracellular delivery), which could activate CREB (by S133 phosphorylation) in neurons both *in vitro* and *in vivo* following intraperitoneal (i.p.) administration [36, 41]. TT221 was further shortened to an 11 aa peptide linked to Tat, denoted TT231. As cationic cell-penetrating sequences, such as Tat, have previously shown issues with cytotoxicity [42, 43], we sought to remove the Tat-sequence but preserve the peptides’ capability of activating CREB *in vivo* following intravenous (i.v.) injection (see optimization strategy in Fig. 1A). We designed and tested 28 peptide analogues, all lacking the Tat-sequence, for their ability to activate CREB (S133) in primary cortical neurons. Here we identified TT241, an 11 aa head-to-tail backbone cyclized peptide, displaying higher efficacy compared to TT231 at 1 µM (Sup. Fig. 1A-B). In addition, TT241 (13 mg/kg) induced a significantly stronger increase in phospho-CREB (S133) levels in the striatum of wild-type mice following i.p. injections compared to 3 other selected peptides of interest (Sup. Fig. 1C-D).

**Fig. 1:**
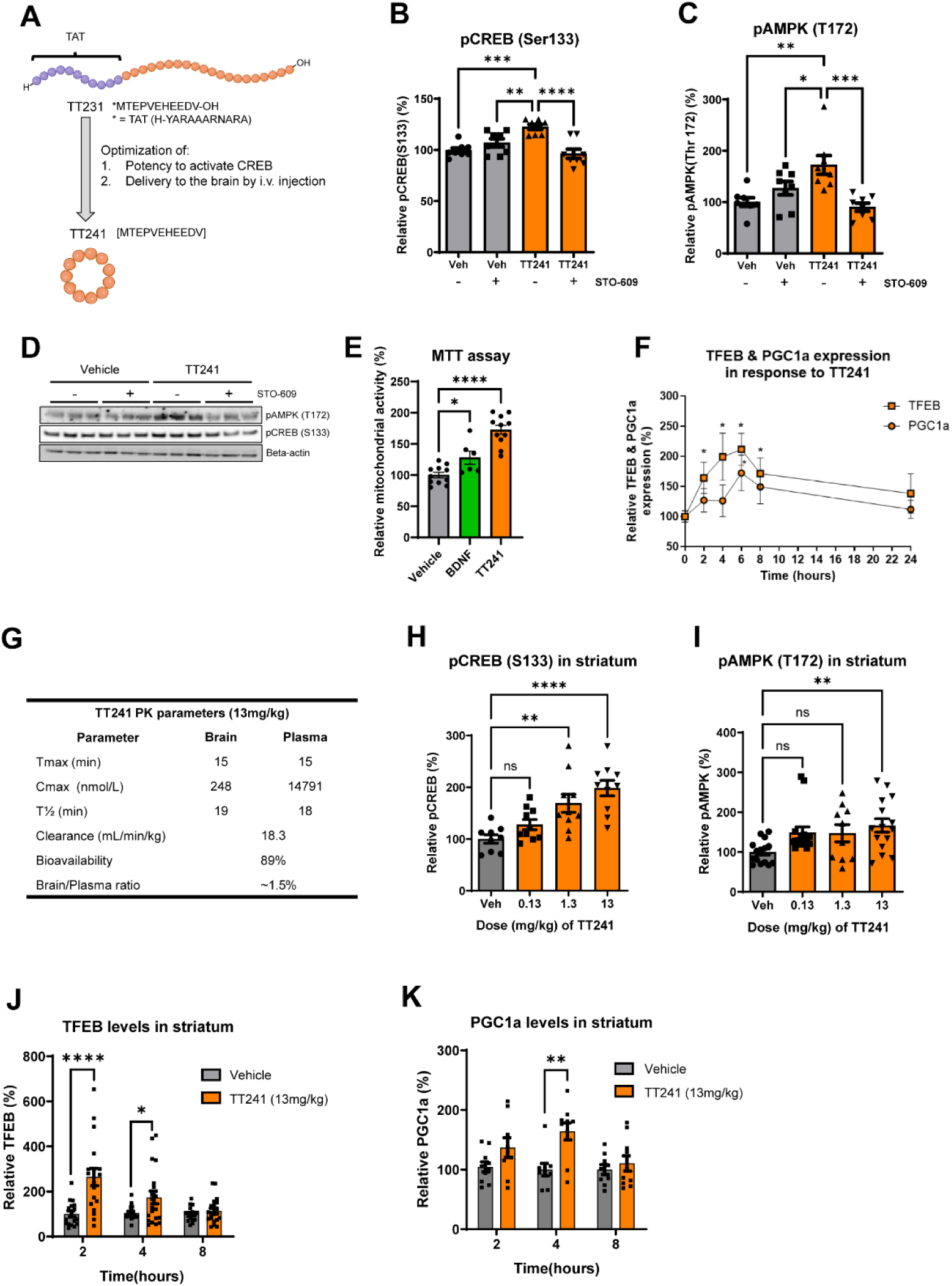
TT241 activates pro-survival signals to induce mitochondrial and lysosomal biogenesis. **(A)** Illustration of the strategy by which TT231 was optimized leading to the cyclic peptide, TT241. **(B-C)** Densitometric analysis of phospho-CREB (S133) and phospho-AMPK (T172) in primary neurons (n=8) with and without TT241 and STO-699 (inhibitor of CAMKK). **(D)** Representative images of phospho-CREB, phospho-AMPK and beta-actin. **(E)** MTT viability assay in primary cortical neurons treated with BDNF or TT241 till 12DIV (n=6-11). **(F)** TFEB and PGC1a time-dependent expression following TT241 treatment in primary cortical neurons (n=8). **(G)** Pharmacokinetics parameters of TT241 in C57BL/6J mice. **(H-I)** Relative dose-response effect of TT241 on phospho-CREB (S133) and phospho-AMPK (T172) in the striatum of C57BL/6J mice when administered intravenously (i.v.) (n=10-15). **(J-K)** Relative time-dependent effect of TT241 on PGC1A and TFEB in the striatum of C57BL/6J mice when administered i.v. (n=10-20). Significance was calculated using ordinary one-way ANOVA. Mean ± SEM.

SorCS2 has been shown as an indispensable regulator of N-methyl-D-aspartate receptor (NMDAR)-dependent LTP formation [26], which requires *de novo* protein synthesis through Calcium/calmodulin-dependent protein kinase IV (CaMKIV)-CREB pathway [44, 45]. To further elucidate the mechanism by which TT241 activates CREB, we evaluated whether this was mediated in a CaMKIV-dependent manner. To assess this, we treated primary neurons with TT241 (1 µM) together with the CaMKK2-selective inhibitor STO-609 (5 µM), which is the upstream CaM-dependent kinase activator of CaMKIV [46–48]. In addition, we assessed the phosphorylation (T172) of AMP-activated protein kinase, AMPK, and CREB which are both targets of CaMKK2, to understand the pathway specificity. Interestingly, we found that TT241 activated both AMPK (T172) and CREB (S133), and this effect was lost in the presence of STO-609, suggesting that TT241 activates CREB and AMPK through CaMKK2 (Fig. 1B-D).

AMPK is an important mediator of synaptic plasticity during synaptic activation as it maintains mitochondrial respiration when energy demands are high [49]. CREB has likewise been linked to metabolic activity and bioenergetics in neurons through regulation of mitochondrial gene expression [50, 51]. We therefore assessed whether TT241 increases metabolic activity in primary neurons using MTT [3-(4,5-dimethylthiazol-2-yl)-2,5-diphenyltetrazolium bromide) assay over an extended treatment period. Indeed, treatment of primary neurons with TT241 (1 µM) led to a significant increase in metabolic activity after 5 days of treatment (Fig. 1E), showing that TT241 targets metabolic pathways to increase metabolic activity. This is in line with previous data on peptide TT221, which induced expression of genes involved with oxidative phosphorylation[36]. Both AMPK and CREB have been shown to regulate mitochondrial biogenesis through upregulation of the mitochondrial master regulator Peroxisome proliferator-activated receptor gamma coactivator 1-alpha (PGC1α), as a part of the CaMKK2-CaMKIV signaling pathway [52–54]. Of note, the expression and activity of PGC1α is tightly interlinked with the lysosomal master regulator Transcription factor EB (TFEB), which is similarly regulated by CREB and AMPK [55–57]. Thus, we speculated that TT241 might increase mitochondrial activity through regulation of PGC1α and TFEB. We therefore assessed expression of both master regulators following acute stimuli with TT241 (1 µM). Both PGC1α and TFEB showed a time-dependent increase, which was significant for TFEB at 2-8 hours and significant for PGC1α at 6 hours (Fig. 1F). We next assessed whether these effects could be obtained *in vivo* in the brain following administration of TT241. Initially, the exposure levels of TT241 in plasma and brain were evaluated following i.v. or s.c. administration of 13 mg/kg. Both i.v. and s.c. administration showed similar exposure profiles in plasma and brain (Sup. Fig. 1E-F) indicating TT241 can be administered to the brain by s.c. administration with a half-life of ∼20 minutes. The pharmacokinetic properties of TT241 following s.c. administration in mice are shown in Fig. 1G and TT241 plasma protein binding in Sup. Fig. 1G. In the following *in vivo* dose-response study, a significant dose-dependent effect on striatal phospho-CREB (S133) was observed (1.3 mg/kg and 13 mg/kg) following s.c. administration; whereas, phospho-AMPK (T172) was only significantly increased at the highest dose of 13 mg/kg (Fig. 1H-I). The activation of AMPK and CREB translated into a time-dependent increase in striatal TFEB and PGC1α. The expression curves were similar to the curves observed *in vitro*, as TFEB was rapidly induced after 2 hours and PGC1α was only significantly increased 4 hours after administration (13 mg/kg) (Fig. 1J-K). Together these data support the ability of TT241 to engage in mitochondrial pathways of the striatum following s.c. administration.

### A stabilized and lipidated peptide analogue, TT-P34, prolongs pathway activation

Next, we asked whether we could prolong the activity of SorCS2 derived peptides *in vivo* by reducing its clearance. TT241 displayed a half-life of ∼20 min (Fig. 1G) and 7.5 % plasma protein binding in mice (Sup. Fig. 1G). We analyzed 40 new peptide analogues for plasma protein binding, half-life, metabolic stability and delivery to the brain after s.c. administration, which yielded an optimized peptide, TT-P34 (Fig. 2A, data for other peptides not shown). TT-P34 is an 11 aa head-to-tail backbone cyclized peptide conjugated to a C_18_ diacid lipid through γGlu and 2 x OEG (amino-3,6-dioxaoctanoic acid) linkers (C_18_DA-yGlu-OEG-OEG). This lipid had previously been used in the development of GLP-1 agonists such as semaglutide [58, 59]. TT-P34 further constitutes two amino acid changes at an identified cleavage site (by *in vitro* metabolite identification study in mouse brain homogenate) in the TT241 peptide sequence (Pro4Gln; Val5Ile).

**Fig. 2:**
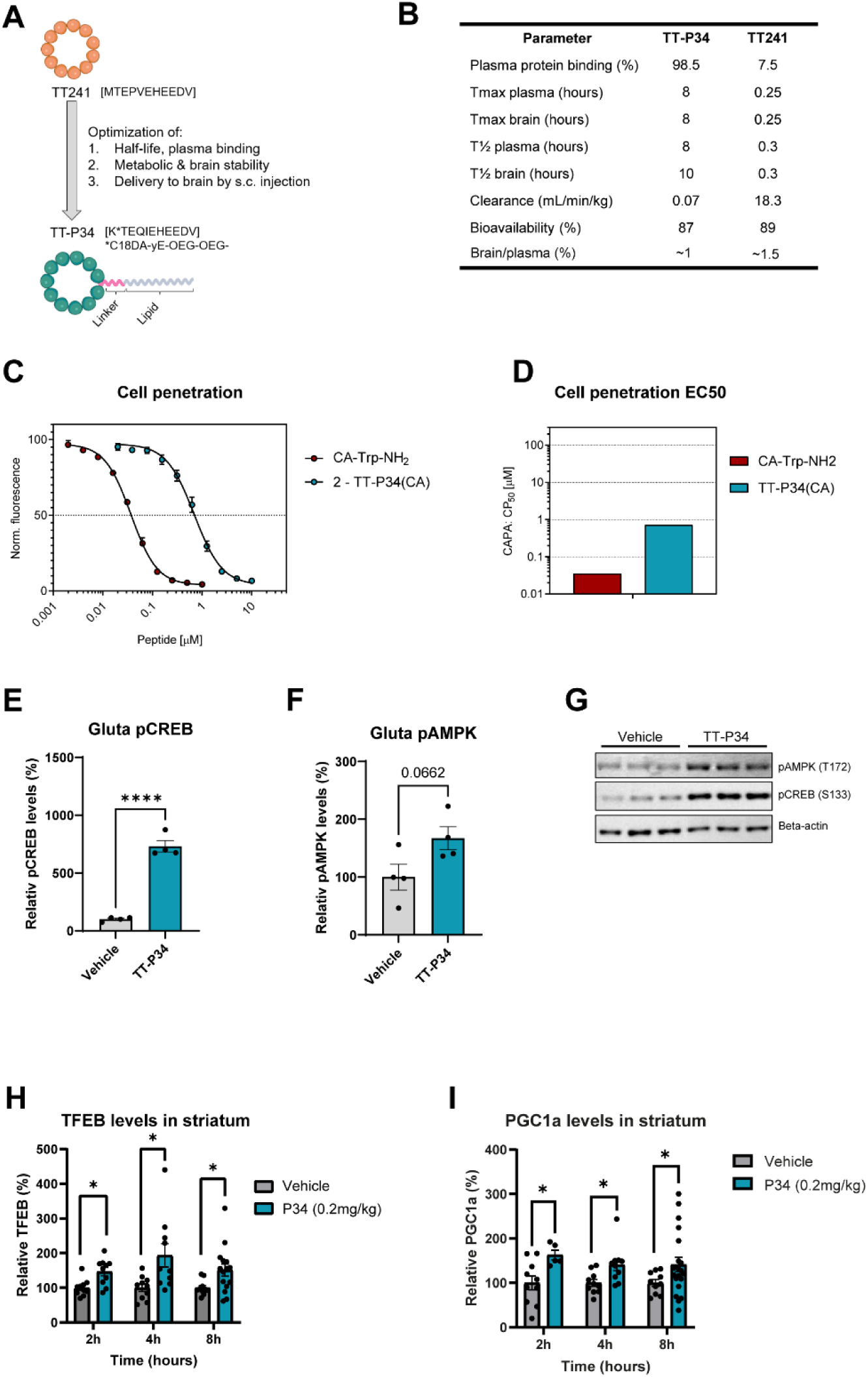
TT-P34 an optimized peptide displays prolonged half-life and extended signaling. **(A)** Illustration of the strategy by which TT241 was optimized leading to the lipidated cyclic peptide, TT-P34. **(B)** Comparison of the pharmacokinetic profiles of TT-P34 and TT241 in C57BL/6J mice. **(C)** Dose-response curves of small molecule Trp and TT-P34 peptides tagged with chloroalkane in chloroalkane penetration assay (CAPA) in Hela cells. **(D)** Cell-permeability (CP50) concentrations obtained in CAPA assay. **(E-F)** Densitometric analysis of phospho-CREB (S133) and phospho-AMPK (T172) in human iPSC-derived glutamatergic neurons (n=4) with and without TT-P34. **(G)** Representative images of phospho-CREB, phospho-AMPK and beta-actin. **(H-I)** Relative time-dependent effect of TT-P34 on PGC1A and TFEB in the striatum of C57BL/6J mice when administered subcutaneously (s.c.) (n=10-15). Significance was calculated using Student’s two-tailed t-test at every timepoint. Mean ± SEM.

TT-P34 demonstrated increased plasma protein binding in mouse plasma with approximately 99% bound, which was the same for rat, dog, human and non-human primate (Sup. Fig. 2A). Furthermore, TT-P34 had superior brain stability in mouse brain homogenates as it was stable for 8 hours of incubation (Sup. Fig. 2B). This translated into increased half-life in mouse brain and plasma following s.c. administration (Sup. Fig. 2C). A comparison of the pharmacokinetic properties of TT241 and TT-P34 is shown in Fig. 2B. Given that TT-P34 is derived from the intracellular domain of SorCS2, which resides in the cytoplasm, we further analyzed the cell-permeability of chloroalkane-tagged TT-P34 using the chloroalkane penetration assay (CAPA) in HeLa cells that allows direct estimation of the cytosolic penetration of molecules [60–63]. We observed cell-permeability and thus cytosolic delivery of TT-P34 with an estimated CP_50_ of 0.72 µM (Fig. 2C-D). Instability issues in lysate were likewise evaluated, in which TT-P34 was completely stable (Sup. Fig. 2D-E). Subsequently, we assessed TT-P34 activity in human induced pluripotent stem cell (iPSC)-derived neurons. Here, TT-P34 potently activated both CREB (S133) and AMPK (T172) at 10 nM (Fig. 2E-G) in human iPSC-derived glutamatergic neurons. This effect was also evident in human iPSC-derived gamma-aminobutyric acid expressing (GABAergic) neurons (Sup. Fig. 2F-H), demonstrating that TT-P34 is biologically active in both excitatory and inhibitory neurons.

To understand the *in vivo* pharmacodynamics, we administered several doses (0.002-2 mg/kg) of TT-P34 and analyzed striatal TFEB levels after 2 hours of exposure in mice. TT-P34 significantly increased TFEB levels in the striatum at 0.2 mg/kg (Sup. Fig. 3A-B) - an approximately 100x lower dose than the E_max_ for TT241 (13 mg/kg) supporting improved potency. Finally, to understand whether the extended half-life translated into a prolonged activity profile, we analyzed the time-dependent increase of TFEB and PGC1α in the striatum of mice following s.c. administration of 0.2 mg/kg. Indeed, TT-P34 displayed a changed pharmacodynamic profile, compared to TT241, as the induction of both TFEB and PGC1α were prolonged with significantly increased levels up to 8 hours following s.c. administration (Fig. 2H-I), which was not evident for TT241 (Fig. 1J-K). In addition, TT-P34 significantly increased the neurotrophic factor BDNF, a transcriptional target of CREB in BDNF/Tropomyosin receptor kinase B (TrKB) signaling [64], at 4 hours post administration (Sup. Fig. 3C-D). Taken together, we developed TT-P34, a lipidated macrocyclic peptide, with prolonged and more potent *in vivo* activity compared to TT241, through optimization of half-life and metabolic stability.

### TT-P34 rescues the behavioral phenotype in zQ175 mouse model of Huntington’s

One of the hallmarks of Huntington’s Disease (HD) is the mutant huntingtin (mHTT) mediated impairment of BDNF-signaling and reduced transcriptional activity of CREB in medium spiny neurons of the striatum, which ultimately results in neuronal death [13, 65, 66]. As the SorCS2-derived peptides activated CREB and in consequence target key disease mechanisms of HD, we next evaluated the therapeutic potential of TT241 and TT-P34 in models of this disease. The PGC1α/TFEB pathway has previously shown to clear Huntington’s disease-causing aggregates of mHTT [67] and thus we initially assessed whether TT241 and TT-P34 could reduce mHTT in patient-derived fibroblasts, based on their effects on PGC1α/TFEB pathway activation. To evaluate this, we treated patient-fibroblasts for 3 days with vehicle, TT241 or TT-P34 and measured both mHTT and total HTT by western blotting. We observed a significant reduction of 25 % of mHTT while total HTT levels remained unchanged (Fig. 3A-C). This supports that TT241 and TT-P34 can activate their target pathways in the presence of mHTT.

**Fig. 3:**
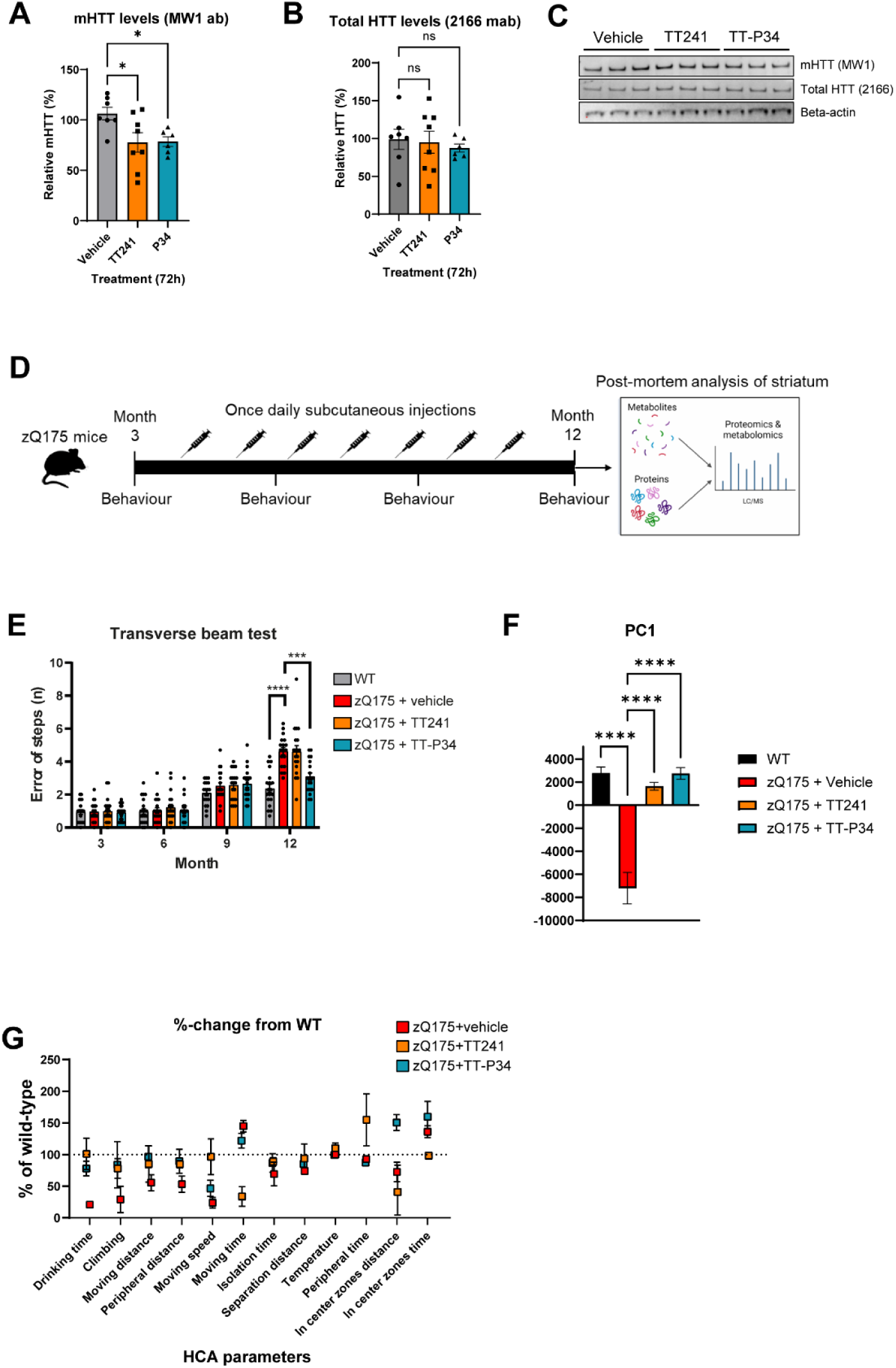
TT-P34 rescue motor functions in zQ175 mouse model of Huntington’s. **(A)** Densitometric analysis of mutant Huntingtin (mHTT) and **(B)** total Huntingtin (HTT) levels in Huntington’s patient-derived fibroblasts treated with TT241 or TT-P34 for 72 hours (n=6-8). **(C)** Representative images of mHTT, total HTT and beta-actin. **(D)** Illustration of zQ175 experiment setup with treatment regime, timepoints with behavioral tests and post-mortem analysis. **(E)** Number of error of steps in the transverse beam test at 3-12 months of age (n=18-20). **(F)** Principal component 1 from principal component analysis (PCA) of home-cage analysis (HCA) data at 12-months of age. **(G)** Relative changes in each behavioral parameter measured in the HCA compared to wild-type (n=8). Significance was calculated using ordinary one-way ANOVA. Mean ± SEM.

As TT-P34 and TT241 exhibited comparable *in vitro* efficacies in terms of mHTT lowering but their pharmacokinetic properties are different *in vivo*, we wanted to test their effects in an *in vivo* model of Huntington’s Disease. Accordingly, TT241 and TT-P34 were evaluated in the zQ175 mouse model of HD, which has the mouse Htt exon 1 replaced by the human Htt exon 1 with ∼190 CAG repeats and is a well-established model to study mechanisms and therapeutic interventions of HD [68, 69].

We treated zQ175 heterozygous mice by once-daily s.c. administration with either vehicle, TT-P34 (0.2 mg/kg) or TT241 (13 mg/kg) from 3 to 12 months of age. Wild-type (WT) littermates were included as controls. Throughout the study, the animals were measured on motor and behavioral functions by rotarod and transverse beam test. In addition, activity was measured by home cage analysis (HCA) using ActualHCA™, which includes analysis of 12 behavioral parameters of social and motor activity, among others. At 12 months of age, the mice were MRI scanned followed by terminal striatal tissue collection for metabolomics and proteomics analysis. An illustration of the setup is shown in Fig. 3D. We observed that neither TT241 nor TT-P34 had any effect on body-weight throughout the study, while all zQ175 heterozygous mice showed significantly reduced body-weights compared to WT (Sup. Fig. 4A). No behavioral changes were observed in rotarod throughout the study (Sup. Fig. 4B). Notably, the vehicle-treated zQ175 mice displayed a significantly increased number in error of steps at 12 months of age compared to WT, which was fully rescued by TT-P34 treatment but not with TT241 treatment (Fig. 3E). To analyze the data generated by HCA, we used a principal component analysis (PCA) of all the behavioral parameters to classify the groups. Here, at 12 months of age, the vehicle-treated zQ175 mice showed significant behavioral changes compared to WT on principal component 1 (explaining 98.2% of all the variance, Sup. Fig. 4C) shown in Fig. 3F. Notably, treatment with TT-P34 completely rescued the behavioral phenotype; TT241 treatment also had a significant effect (Fig. 3F). A PCA plot of PC1 and PC2 (capturing all the variance) shows the same trend, i.e. that TT-P34-treated zQ175 mice behave almost identical to WT (Sup. Fig. 4D). Breaking down each of the parameters relative to WT levels revealed that both TT-P34 and TT241 rescue motoric deficits involving climbing, moving distance, moving speed and peripheral distance while also restoring drinking time (Fig. 3G). Surprisingly, the behavioral changes were not coupled with any changes in brain volume measured by MRI, which were significantly lower in all zQ175 mice compared to WT (Sup. Fig. 4E). Taken together, the data shows that the SorCS2-derived peptides rescued the behavioral phenotype of the zQ175 mouse model of HD and further supports that sustained activity by TT-P34 was more efficacious than a transient activity displayed by TT241 *in vivo*.

### TT-P34 increases proteins of oxidative phosphorylation and prevents loss of synaptic content

While we showed that the SorCS2-derived peptides TT241 and TT-P34 engage with the PGC1α and TFEB pathways we were unaware of the broader molecular consequences of sustained activation of this system *in vivo*. Thus, we conducted an unbiased analysis of the striatal tissue by subjecting it to LC-MS based label-free proteomics and metabolomics analysis testing the TT-P34-treated group. We identified significant changes with TT-P34 treatment constituting a total of 283 increased and 38 decreased proteins as well as 123 increased and 51 decreased metabolites (Fig. 4A-B and Sup. Fig. 5A-B). Integrated pathway analysis of all significant changed proteins and metabolites with TT-P34 treatment identified enriched pathways of neurodegenerative diseases including PD, AD and HD and pathways of both the dopaminergic synapse, retrograde endocannabinoid signaling as well as oxidative phosphorylation among others (Fig. 4C). Notably, the proteins significantly changed by TT-P34 treatment annotated to the oxidative phosphorylation pathway were almost exclusively NADH:ubiquinone oxidoreductase (*Nduf*) subunits, which are part of mitochondrial complex 1 (MC1) (Fig. 4D). Cytochrome C1 (*Cyc1*) and Cytochrome c oxidase 6B1 (*Cox6b1*) were the only ones not part of MC1. In line with this, NAD^+^ levels were significantly elevated with TT-P34 treatment, although not significantly downregulated in the zQ175 model compared to WT (Fig. 4E). This clearly demonstrated that TT-P34 increased metabolic activity potentially as a consequence of TFEB and PGC1α pathway engagement. As TFEB is a driver of lysosomal biogenesis, we additionally explored the significance of sustained TT-P34 treatment on lysosomal proteins. A few lysosomal proteins including Rabs (Rab3c, Rab18, Rab5a, Rab7) and V-type proton ATPase catalytic subunits (Atp6v1g1, Atp6v1a, Atp6v0d1) were significantly changed with TT-P34 treatment (Sup. Fig. 5C) supporting the regulation of the lysosomal network.

**Fig. 4:**
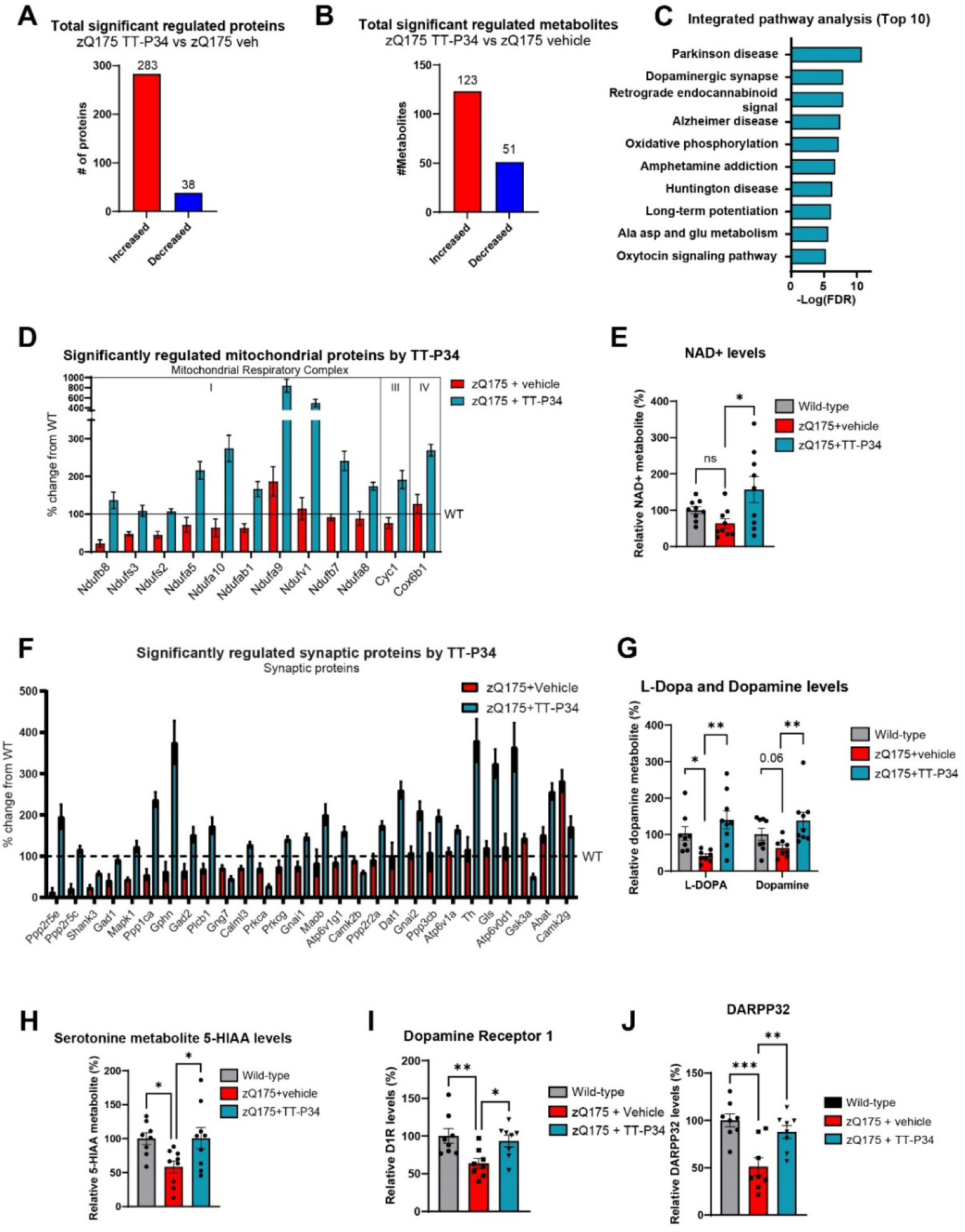
TT-P34 increase mitochondrial proteins and rescues loss of synaptic content in striatum. **(A)** Total significant changed proteins and **(B)** metabolites in striatum of zQ175 treated with TT-P34 (FDR<0.05) identified by proteomics and metabolomics. **(C)** Enriched pathways found using integrated pathway analysis of all significantly changed proteins and metabolites with TT-P34 treatment. **(D)** Significant changed striatal proteins with TT-P34 treatment (TT-P34 treated vs zQ175 vehicle treated) annotated to the oxidative phosphorylation pathway taken relative to wild-type levels (n=9). **(E)** Relative NAD+ metabolite levels in the striatum of zQ175 treated with vehicle or TT-P34 compared to wild-type (n=9). **(F)** Significant changed striatal proteins with TT-P34 treatment (TT-P34 treated vs zQ175 vehicle treated) involved in synaptic functions taken relative to wild-type levels (n=9). **(G)** Relative striatal L-dopa, dopamine and **(H)** 5-HIAA serotonin metabolites of zQ175 mice compared to wild-type (n=9). **(I-J)** Relative levels of medium spiny neuron markers D1R and DARPP32 measured by simple western (n=8). Significance was calculated using ordinary one-way ANOVA. Mean ± SEM.

The pathway enrichment analysis revealed a significant change in synaptic pathways with treatment, where almost all proteins annotated hereto were upregulated (Fig. 4F). Of note, proteins found primarily at glutamatergic synapses (Gls, Shank3), dopaminergic synapses (Th, Dat1, Maob) and GABAergic synapses (Gphn, Gad1, Gad2, Abat) were all significantly increased suggesting that TT-P34 treatment affects multiple neuronal systems. The synaptic engagement was also evident from a significant rescue of L-3,4-dihydroxyphenylalanine (L-DOPA), dopamine and the serotonin metabolite, 5-HIAA (Fig. 4G-H) and significant increases in L-glutamic acid and γ-aminobutyric acid (GABA) (Sup. Fig. 6A-B). The broad positive effects on mitochondrial and synaptic proteins and metabolites also translated into a rescue of medium spiny neuronal markers of dopamine receptor 1, DARPP32 and SCN4B validated by simple western (Fig. 4 I-J & Sup. Fig. 6C). This was accompanied with significant increases in both PGC1α and TFEB levels, although these were not themselves downregulated in the zQ175 model (Sup. Fig. 6D-E). Finally, we measured neurofilament light chain levels in both plasma and CSF as a well described biofluid biomarker of disease progression in HD, where a trend (but no significant effect) appeared with treatment (Fig. 6F-G). Taken together, this shows that TT-P34 rescues synaptic proteins and metabolites in the zQ175 model potentially by regulating mitochondrial function through PGC1α/TFEB regulation.

### TT-P34 ameliorates the behavioral phenotype and rescues dopaminergic terminals in a MPTP mouse model

Mitochondrial deficits are a key pathogenic mechanism in PD, which was originally discovered by patients being exposed to the mitochondrial toxin 1-methyl-4-phenyl-1,2,3,6-tetrahydropyridine (MPTP; precursor to neurotoxin MPP^+^) causing permanent parkinsonian-like symptoms through inhibition of mitochondrial complex I (MCI) in dopaminergic neurons and subsequent neuronal degeneration [70, 71]. Based on this, animal models using MPTP have been extensively characterized and used as models for PD. As both TFEB and PGC1α overexpression have previously shown beneficial effects in these models, we evaluated TT-P34 in an MPTP mouse model of PD [72–74].

TT-P34 was administered s.c. once-per day at five different doses (0.0016-1 mg/kg) to wild type mice, starting one-day prior to treatment with MPTP (30 mg/kg/day). Baseline and endpoint behaviors including open field test (OFT), grip strength and body weight were measured. Tyrosine hydroxylase (TH) was stained by immunohistochemistry to evaluate dopaminergic survival in substantia nigra pars compacta (SNpc). A schematic overview of the experiment is shown in Fig. 5A.

**Fig. 5:**
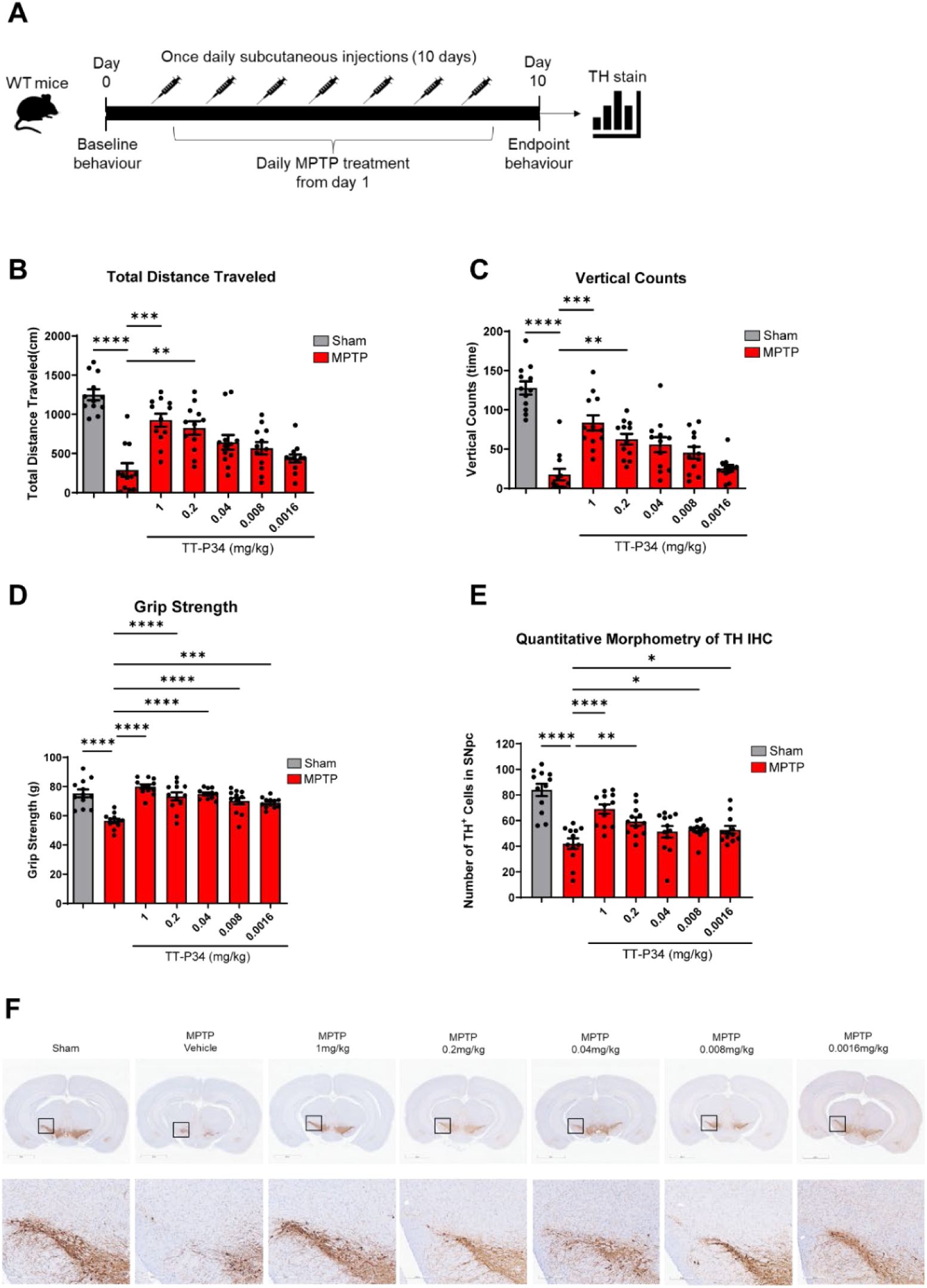
TT-P34 ameliorates motor dysfunctions and rescue dopaminergic neurons in MPTP-mouse model. **(A)** Illustration of MPTP-induced parkinsonism experiment setup with treatment regime, timepoints with behavioral tests and post-mortem analysis. **(B)** Endpoint distance travelled(cm) and **(C)** vertical counts measured in open-field analysis of mice administered with vehicle (sham) or MPTP +-different doses of TT-P34 for 10 days by subcutaneous injections (n=12). **(D)** Grip strength measurements at endpoint (n=12). Kruskal-Wallis test was used for statistical significance. **(E)** Quantitative morphometry of tyrosine-hydroxylase (TH) stainings in substantia nigra and **(F)** representative images of TH-stains for each group with sham-treated (no MPTP) and MPTP-treated with the different doses of TT-P34 displayed. Ordinary ANOVA was used for statistical significance. Mean ± SEM.

Subchronic dosing across 9 days with MPTP (30 mg/kg/day) resulted in a small drop in body weight which TT-P34 did not affect (Sup. Fig. 7A). In the OFT, vehicle treated MPTP mice displayed a significant reduction in total distance travelled and vertical activity compared to sham treatment (did not receive MPTP). Notably, TT-P34 showed a clear dose-response effect in both OFT measurements with a significant ameliorative effect at the highest doses of 0.2 and 1 mg/kg (Fig. 5B-C, p<0.05, Kruskal-Wallis test). Additionally, the grip-strength was significantly rescued across all doses (Fig. 5D). Post-mortem staining and quantitative morphometry analysis of TH-positive cells in SNpc demonstrated a 50% loss in the MPTP-vehicle treated group compared to the sham-treated group (Fig. 5E-F). The TH-positive neurons were significantly rescued to ∼90% of normal levels with 1 mg/kg dosing and ∼70% of normal levels with 0.2 mg/kg. Doses of 0.008-0.0016 mg/kg showed effects, although more modest. Further validation of TT-P34 levels in plasma demonstrated a linear relationship between doses, at which 1 mg/kg led to an estimated free brain concentration of 12 ng/mL, corresponding to 6 nM of free TT-P34 (Sup. Fig. 7B-C). Together, this demonstrates that TT-P34 rescues dopaminergic loss and restores behavioral deficits caused by MPTP-mediated MCI inhibition with the best effect obtained at 6 nM free brain concentrations after 10 days of dosing.

### A human dose prediction for TT-P34

Given the clear therapeutic benefit of TT-P34 in two models of NDDs, we next sought to explore the pharmacokinetic and pharmacodynamic profiles of TT-P34 in larger animals to better understand its translatability and to inform on a putative human dosing regimen. To do so, we designed and carried out four studies which included 1) evaluation of pharmacokinetics (PK) in mice, rats, dogs and non-human primates (NHPs) to predict human PK; 2) understanding the pharmacodynamics on TFEB as a brain biomarker of efficacy; 3) coupling TFEB brain biomarker efficacy with therapeutic dose efficacy in the MPTP mouse model; and 4) establishing a physiologically based pharmacokinetic model (PBPK) for predicting brain penetration in humans. A schematic of the process is shown in Fig. 6A. In combination, the four data packages would help give confidence in the proposed human dosing regimen.

**Fig. 6:**
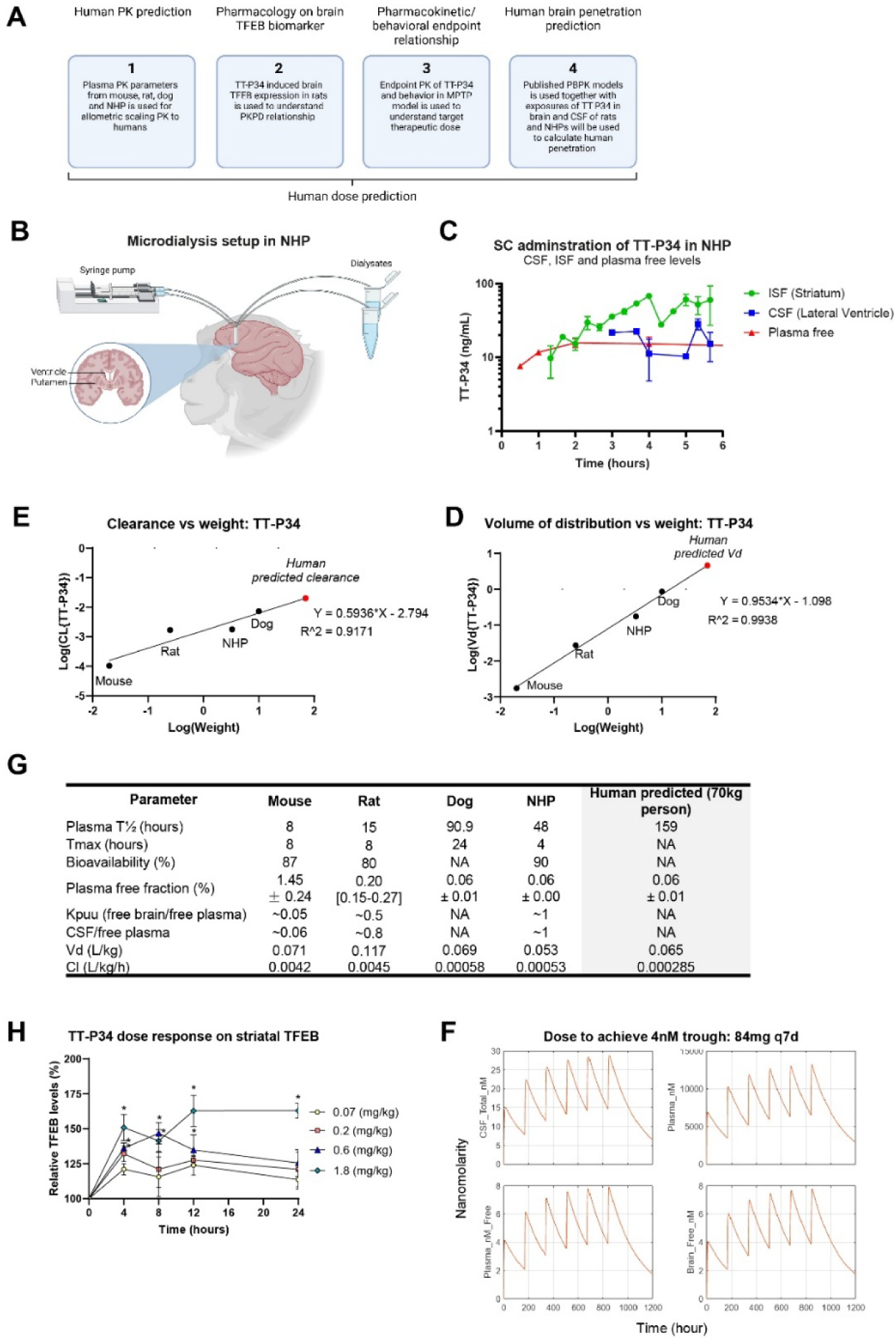
Human dose prediction for TT-P34. **(A)** Acute dose-response effect of TT-P34 on TFEB expression in striatum over time when administered subcutaneously (s.c.) in Sprague Dawley rats (n=5). Ordinary ANOVA was used for statistical significance. Mean ± SEM. Modelling the response gives rise to an indirect PKPD model, where a separate effect site (brain) occurs outside of the central compartment (blood) and the drug distribution to that compartment causes the delay of effect. The model parameters are shown in **(B)** with a synthesis rate (Ksynth), degradation rate (Kdeg) and an EC50 of 2nM plasma free TT-P34 concentration. **(C)** Illustration of microdialysis setup in non-human primates (Macaca fascicularis) to measure brain penetration of TT-P34 and pharmacokinetics shown in **(D)** following s.c. administration (n=3). Mean ± SEM. Plot of obtained **(E)** clearance rates and volume of distributions **(F)** in mouse, rat, non-human primate and dog versus their weight. Linear regression is used to predict the corresponding clearance rate and volume of distribution in humans (noted as red dot) (n=3). **(G)** Pharmacokinetic profile of TT-P34 in different species used to predict human pharmacokinetics. **(H)** Human dose prediction to achieve 4nM brain-free concentrations a trough accompanied with plasma, plasma-free and total CSF levels.

Detailed PK parameters were initially estimated independently in four species: mouse, rat, dog and NHP following s.c. administration at different doses. I.v. administration was also performed in mice, rats and NHPs to obtain bioavailability. Both plasma, CSF and brain were sampled from mice and rats across timepoints. In dogs only plasma was sampled. In NHPs a microdialysis procedure was performed where CSF from the lateral ventricle and the interstitial fluid (ISF) of the striatum were sampled 1-6 hours post dosing (see schematic in Fig. 6B), thereby giving a direct measure of the free brain and CSF concentrations of TT-P34. Interestingly, we observed a 3-fold difference between free brain concentrations (ISF levels) versus CSF levels in NHPs (Fig. 6C), whereas the CSF levels were equal to the estimated plasma free concentrations. This demonstrated that TT-P34 crossed the blood-brain barrier in NHPs following s.c. administration at levels at least equal to the plasma free concentrations. Following s.c. administration in each species, the plasma PK profile was modelled (Sup. Fig. 8A-D) yielding clearance rates (Cl), volume of distribution (V_d_) and the first-order absorption rate constant (k_a_) based on a one-compartment model, which best described the data. All raw data obtained from each species can be found in Sup. Data File S4. Subsequent allometric scaling was employed to extrapolate the animal PK data to predict human pharmacokinetic parameters. Plotting the logarithmic values of Cl rates and V_d_ vs. weight for each species demonstrated a clear linear relationship between weight and pharmacokinetic parameters (Fig. 6E-D). Extrapolation to humans (70 kg person) resulted in a human Cl of 0.000285 (L/kg/h) and V_d_ of 0.065 (L/kg). Using the formula for t_½_ = 0.693 × V_d_ /Cl, the estimated half-life (T_½_) in humans was calculated to be 159 hours (6.6 days). The pharmacokinetic parameters obtained from each species are shown in Fig. 6G.

Next, we evaluated the pharmacodynamics of inducing TFEB expression in striatum of wild-type rats. Here, TT-P34 was dosed at four different concentrations (0.07-1.8 mg/kg) and striatal TFEB levels were evaluated acutely at several timepoints (4-24 hours post injection). Plasma samples were taken at each timepoint. As expected, administration of TT-P34 significantly increased TFEB in a dose-dependent manner (Fig. 6H). A linear relationship between the doses and plasma exposure levels was additionally observed (Sup. Fig. 8E). Modelling the percentage-change response with estimated plasma free-levels of TT-P34 (based on plasma free fraction of rats) we obtained a plasma free EC_50_ of 2 nM (1 nM brain free). The model further predicted synthesis rates (K_synth_) of 1856.6 s^−1^ and degradation rates of (K_deg_) of 19.76 s^−1^ (Sup. Fig. 8F). Combining the PK model for mice with the observed maximal therapeutic efficacy obtained in the MPTP-mouse model at 1 mg/kg dosing, we estimated the plasma-free concentration of TT-P34 for reaching the observed behavioral effects to be 8 nM at trough. Using a K_puu_ value (ratio between free brain and free plasma concentrations) obtained from rat PK studies, this would correspond to an average of 4 nM free brain concentrations. Therefore, therapeutic dose exposure was approximately 4 times the *in vivo* EC_50_ value of TFEB induction which would approach the EC_90_ values and thus, as expected, suggested that a maximal effect on TFEB biomarker is needed to obtain the best disease-modifying effect.

Finally, to predict the human brain penetration and dosing, we utilized a previously published PBPK model originally developed for paracetamol, which accurately predicted the PK in plasma and cerebrospinal fluid (CSF) both in rats and humans [75]. The model was refined to incorporate the PK parameters of TT-P34 obtained from rat and NHP studies. We used the model to predict a weekly human dose (given that t_½_ = 6.6 days) to obtain the disease-modifying effects observed at 4 nM free brain concentrations at trough. Here, the model predicted that 84 mg of TT-P34 given once per week via s.c. administration in humans would result in free brain exposures of 4 nM at trough, thereby yielding an estimate of the human dose regimen (Fig. 6F).

## DISCUSSION

In this study we have developed two novel macrocyclic peptides, derived from a serine motif located in the intracellular domain of the BDNF co-receptor, SorCS2. We demonstrate that they induce AMPK and CREB-mediated mitochondrial programs both *in vitro* and *in vivo* in a CaMKK2-dependent manner. Prolonged activation of this pathway, using the stabilized, lipidated peptide variant, TT-P34, leads to amelioration and rescue of disease behavioral phenotypes in animal models of HD and PD. These findings support a key role of the SorCS2/CaMKK2/AMPK/CREB-pathway in regulating mitochondrial and lysosomal processes and offers a novel approach for protection against neurodegenerative pathologies. Although we demonstrate that peptides derived from the SorCS2 receptor can regulate these pathways, the role of SorCS2 in this context has already been established. Indeed, SorCS2 protects both neurons and pancreatic cells from oxidative stress and loss of SorCS2 leads to significant changes in expression of mitochondrial proteins with NADH dehydrogenases (mitochondrial complex I proteins) as some of the most affected [76, 77]. Consistent with these findings, we noted that TT-P34 treatment in the zQ175 mouse model of HD resulted in elevated levels of striatal NADH dehydrogenases. Notably, these are also the predominant mitochondrial enzymes impacted in PD [78].

Recently, several SNPs in *SORCS2* were identified to be associated with HD [79]. SorCS2 has moreover been functionally linked with HD by mislocalizing in medium spiny neurons of HD patients in addition to displaying reduced striatal immunoreactivity suggesting reduced levels [34]. These data were replicated in the zQ175 model, where loss of SorCS2 further exacerbates motor coordination deficits, supporting a neuroprotective role of the SorCS2 pathway in HD. In the present study, we showed that TT-P34 rescued the disease phenotype in the zQ175 mouse model of HD when administered at an early stage of disease, which was accompanied by a rescue of striatal markers (DARPP32 and D1R) and synaptic content measured by metabolomics and proteomics. This suggests a neuroprotective function of TT-P34 against mHTT dependent striatal degeneration. Indeed, previous studies have shown that activation of both CREB and the PGC1α/TFEB pathway ameliorate behavioral deficits and striatal neurodegeneration in animal models of HD [34, 66, 67]. However, since TT-P34 treatment was started at three months of age in the zQ175 model, an age that aligns with the presymptomatic stages of HD, this does highlight some limitations to this study.

We discovered that TT-P34 regulated the levels of several neurotransmitters in the striatum including dopamine, serotonin metabolite 5-HIAA, glutamic acid and GABA. Of these, the dopamine metabolites were the most affected in the zQ175 model, and TT-P34 completely rescued the levels of both dopamine and its precursor L-3,4-dihydroxyphenylalanine (L-DOPA). Notably, CREB activity increases the expression of TH (the rate limiting enzyme in dopamine production) and loss of SorCS2 has been shown to have an impact on dopamine metabolism and activity [25, 80–82]. In line with this, our proteomics data demonstrate that TH levels are one of the most significantly elevated proteins with TT-P34 treatment in the striatum of zQ175 (Fig. 4F). Today’s medication for HD patients mainly includes symptomatic treatment using dopamine antagonists such as haloperidol and VMAT2 inhibitors such as tetrabenazine to address involuntarily movements (chorea) which is caused by aberrant overactivity of dopamine at synaptic terminals[83]. Dopamine levels have been shown to follow a biphasic progression with early increases causing chorea and then later decreased levels leading to hypokinesia and dystonia [84]. Considering the relationship between TT-P34 and the CREB/TH/dopamine pathway, this suggests that TT-P34 offers potential benefits in the later stages of the disease by increasing dopamine biosynthesis. Importantly, we did not observe any adverse effects on motor behavior in the zQ175 mouse model at early stages of disease (3-6 months of age) corresponding to the hyperdopaminergic stages stating that this mechanism does not aggravate disease phenotype. Together, the potential regulation of the dopamine synthesis and mitochondrial regulation could provide therapeutic benefits of TT-P34 in several neurodegenerative disorders with affection of dopaminergic brain regions.

Finally, we show here that lipidation with C_18_DA-yGlu-OEG-OEG extended half-life, yielding prolonged and superior effects of TT-P34 relative to TT241. By attaching C_18_DA-yGlu-OEG-OEG to our macrocyclic peptide, we obtain a PK profile similar to that of semaglutide across rodents and larger species (dogs and NHPs). While the human PK for TT-P34 still needs to be assessed, including its delivery to the CNS as the immediate first step in the clinic, the use of C_18_DA-yGlu-OEG-OEG therefore offers enhanced half-life and brings more confidence in a human dose regime for TT-P34.

Taken together, we have developed a novel promising macrocyclic peptide, which is administered s.c., crosses the blood-brain barrier and activates neuroprotective pathways in the brain thereby offering a putative novel therapeutic for people suffering from neurodegenerative disorders.

## Acknowledgments

We thank CHDI Foundation for sponsoring the zQ175 animal for this study. We also thank Erwan Bezard (Motac Neuroscience Ltd) for non-human primate work and technical support.

## Data and materials availability

All data are available in the main text or the supplementary materials.

The HD patient fibroblast cell line samples (GM04476) were obtained from the NIGMS Human Genetic Cell Repository at the Coriell Institute for Medical Research under an MTA.

## Author contributions

Conceptualization: AD, MKO, SG, SM

Methodology: AD, MKO, JP, CROB, NB, ER, SLP, LCP, KS, KF, SM

Investigation: AD, MKO, JP, MG, SN, CROB, NB, ER

Funding acquisition: SM

Project administration: AD, MKO, SM

Supervision: SG, SM

Writing – original draft: AD

## List of Supplementary Materials

Supplementary Material and Methods

Fig. S1 to S8

Data file S1 to S4 (Excel files)

