## Supplementary Figures & Methods for "The SorCS2-derived macrocycle TT-P34 drives neuroprotection in animal models of neurodegeneration"

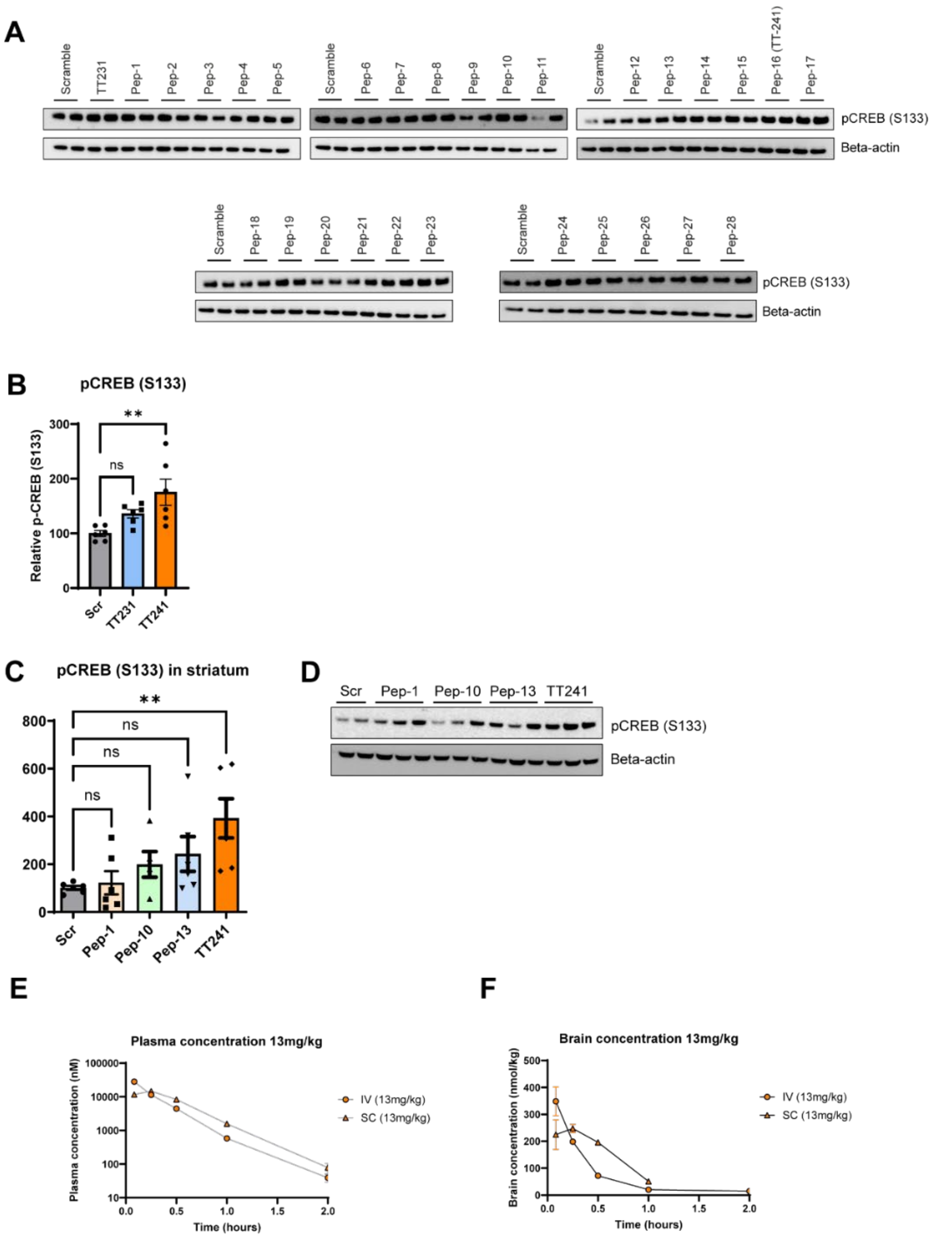


G

| Parameter | Mouse | Human |
|---|---|---|
| Plasma binding (%) | 7.5% | 22% |

**Sup. Fig. 1: Optimization of TT231 and pharmacokinetics of TT241 in wild-type mice.**

**(A)** Representative images of phospho-CREB (S133) and beta-actin induced by peptide analogues of TT231, including TT241 as part of optimization, in primary neurons. **(B)** Densitometric analysis of pCREB normalized to beta-actin (n=6) for TT231 and TT241. **(C)** Densitometric analysis of phospho-CREB (S133) normalized to beta-actin in striatum of wild-type mice following intravenous administration of TT241 and selected peptide analogues of TT231 (n=4-5). **(D)** Representative images of *in vivo* phospho-CREB and beta-actin for selected peptide analogues. Significance was calculated using ordinary one-way ANOVA. **(E)** Plasma and **(F)** brain concentration levels of TT241 following both intravenous (IV) and subcutaneous (SC) administration of 13mg/kg. **(G)** Plasma binding of TT241 in mouse and human plasma samples. Mean ± SEM.

**A**

| TT-P34 plasma free (%) | | | | |
|---|---|---|---|---|
| Mouse | Rat | Dog | Human | NHP |
| 1.45 ± 0.24 | 0.20 [0.15-0.27] | 0.06 ± 0.01 | 0.06 ± 0.01 | 0.06 ± 0.00 |

| TT-P34 brain free (%) | | |
|---|---|---|
| Mouse | Rat | Human |
| 7.4±0.62 | 6.83±0.37 | 11.05 |

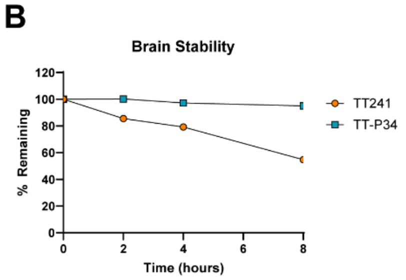


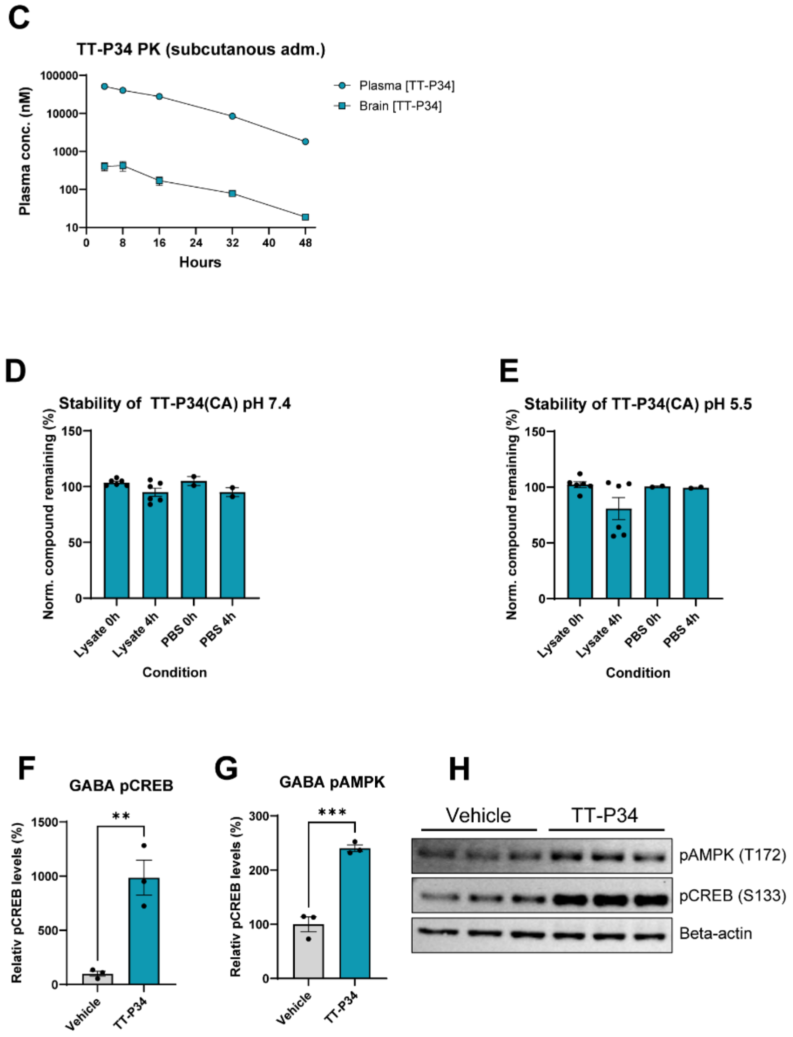


**Sup. Fig. 2: TT-P34 pharmacokinetics and *in vitro* activity in human iPSC-derived GABAergic neurons**

**(A)** Plasma-free and brain-free concentrations of TT-P34 in different species. **(B)** Mouse (C57BL/6J) brain homogenate stability comparison between TT241 and TT-P34. **(C)** Plasma concentration levels of TT-P34 and TT241 following subcutaneous (SC) administration in mice (n=3). **(D)** Stability of chloroalkane-tagged TT-P34 in PBS and HeLa cell lysates at pH 7.4 and **(E)** pH 5.5 after 4 hours (n=6). **(F-G)** Densitometric analysis of phospho-CREB (S133) and phospho-AMPK (T172) normalized to beta-actin in human iPSC-derived GABAergic neurons following TT-P34 treatment (10nM for 15min) and **(H)** representative western blots. Significance was calculated using Student's two-tailed t-test. Mean ± SEM.

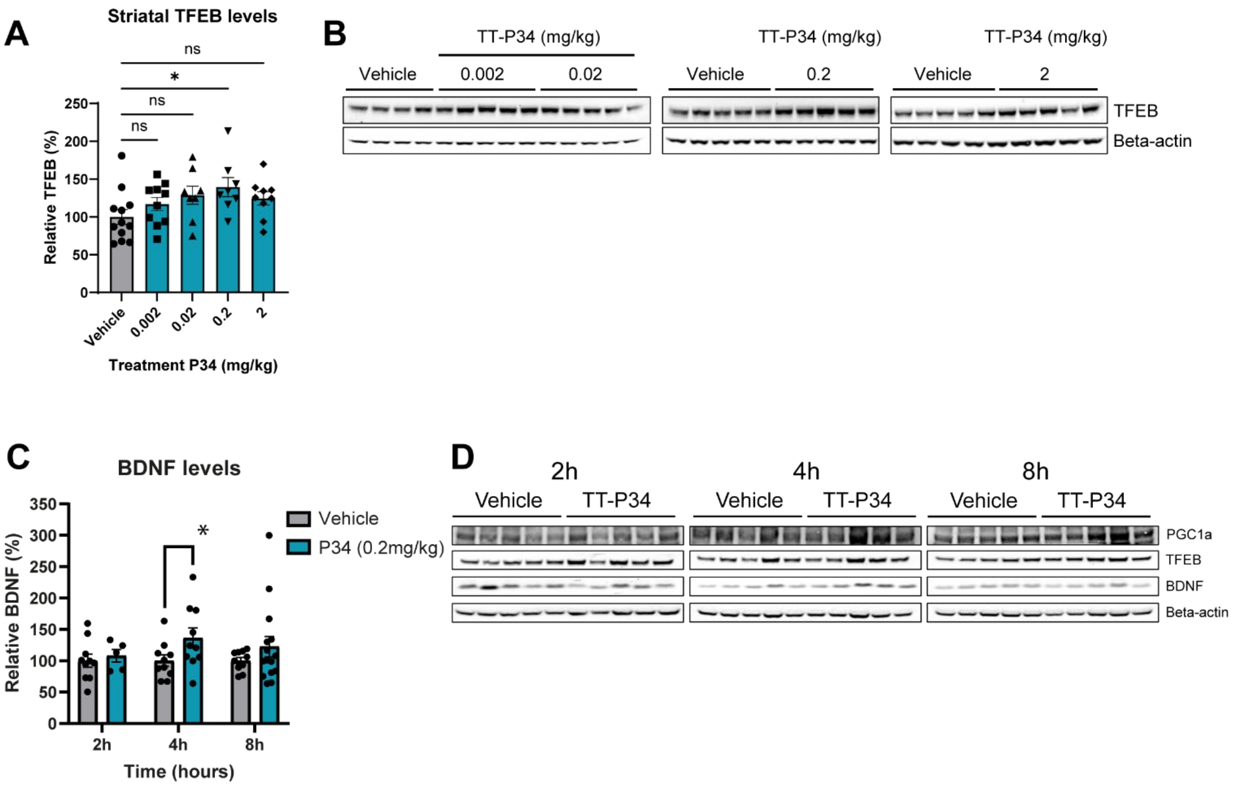


**Sup. Fig. 3: *In vivo* pharmacodynamics of TT-P34**

**(A)** Relative TFEB protein levels normalized to beta-actin in striatum of C57BL/6J following subcutaneous administration of TT-P34 at different doses (n=8-10) and **(B)** representative western blots. Significance was calculated using ordinary one-way ANOVA. **(C)** Relative BDNF protein levels normalized to beta-actin in striatum of C57BL/6J following subcutaneous administration of 0.2mg/kg TT-P34 at 2-8 hours post injection and **(D)** representative images of BDNF, TFEB, PGC1a and beta-actin. Significance was calculated using students two-tailed t-test at each timepoint. Mean ± SEM.

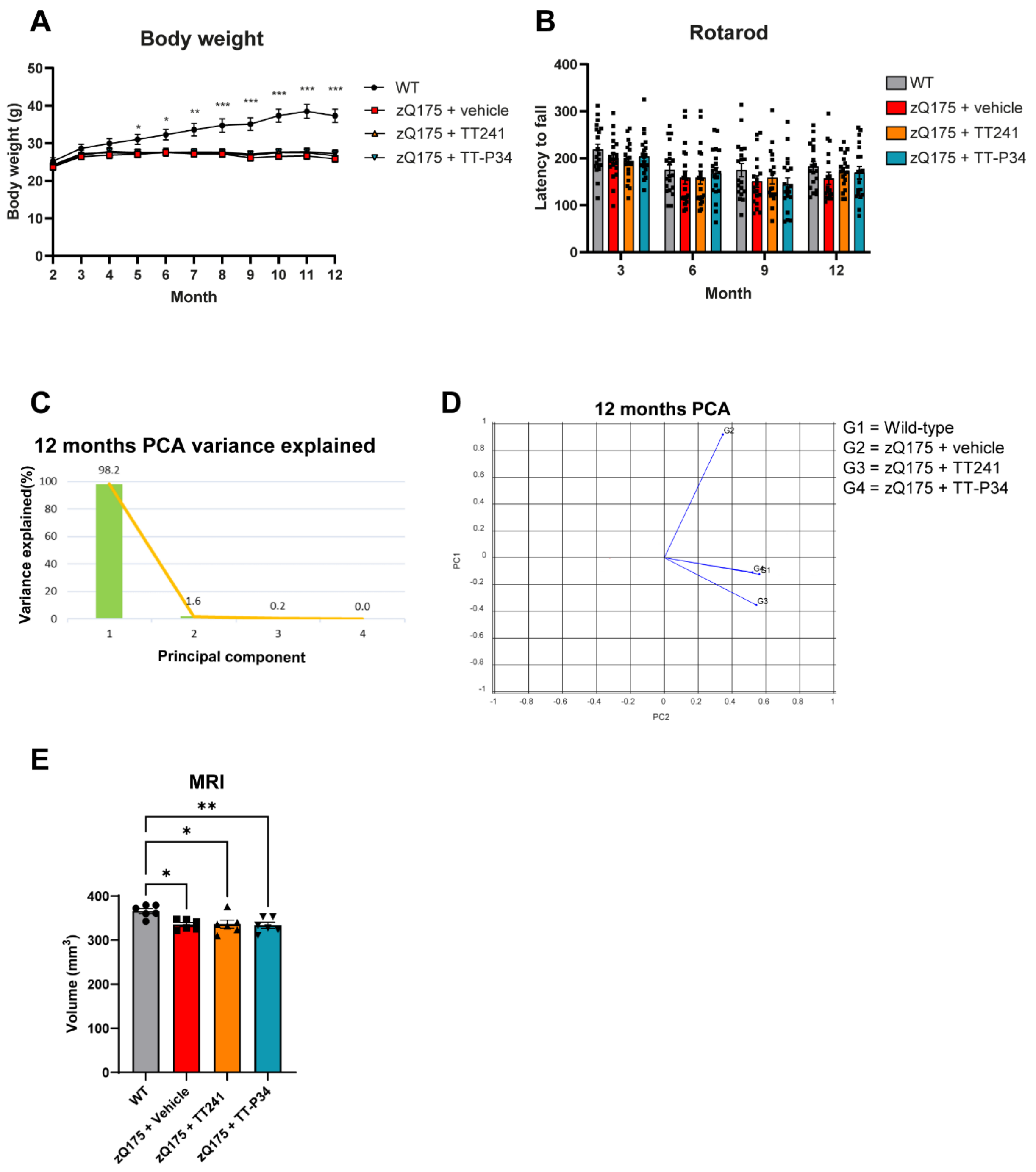


**Sup. Fig. 4: Body-weight, MRI and rotarod in zQ175 mice treated with TT241 and TT-P34**

**(A)** Bodyweight of zQ175 mice treated with once-daily subcutaneous administration of TT-P34 (0.2mg/kg), TT241 (13mg/kg) and vehicle (formulation) and age-matched wild-type (WT) littermates from 3-12 months of age (n=18-20). Significance was calculated using ordinary two-way ANOVA. **(B)** Rotarod at 3, 6, 9 and 12 months of age (n=18-20). **(C)** Principal component variance explained per principal component. **(D)** Principal component analysis plot of PC1 and PC2 from home-cage analysis data at 12 months of age (n=8). **(E)** Whole brain volume measured by MRI of zQ175 mice and WT at 12 months of age (n=6). Significance was calculated using ordinary one-way ANOVA. Mean ± SEM.

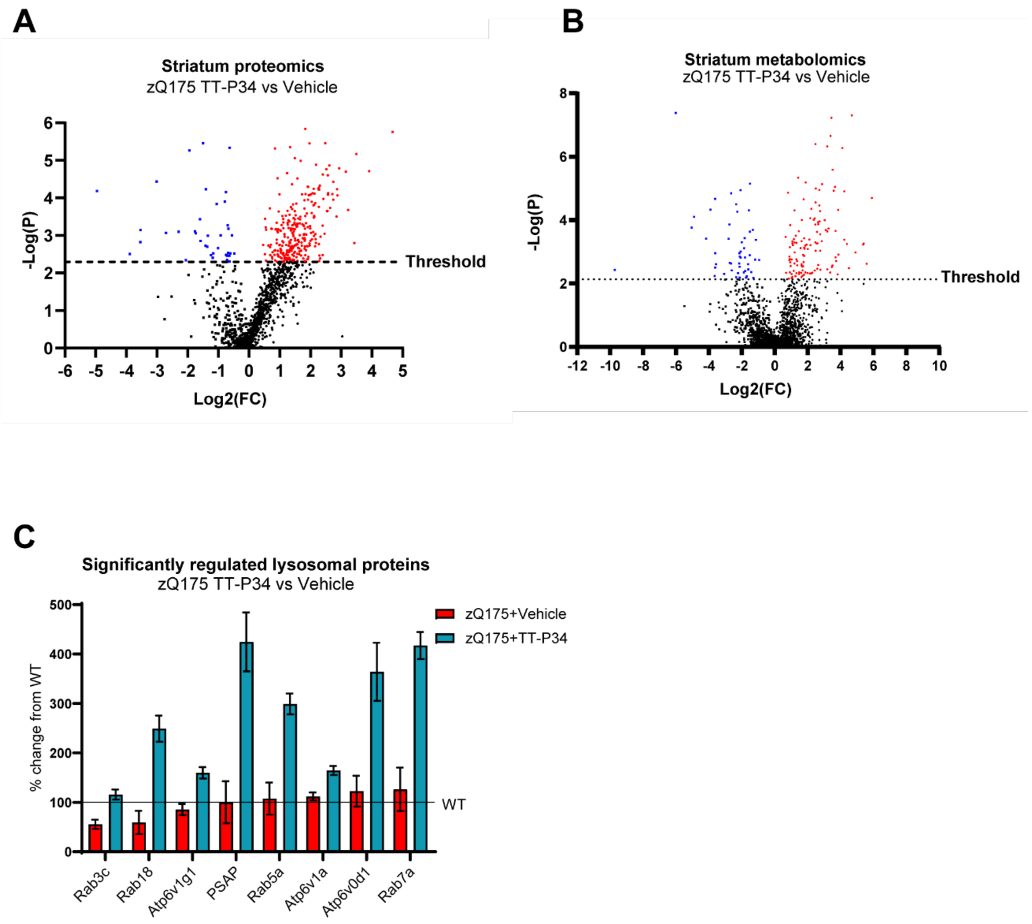


**Sup. Fig. 5: Volcano plot data and lysosomal protein changes from striatum of TT-P34 treated zQ175 mice**

**(A)** Volcano plots of proteomics data and **(B)** metabolomics data of striatal tissue from zQ175 mice treated with TT-P34 compared to vehicle treated (n=9). **(C)** Significantly regulated lysosomal proteins by TT-P34 compared to vehicle-treated normalized to wild-type levels (dotted line) in striatum zQ175 mice.

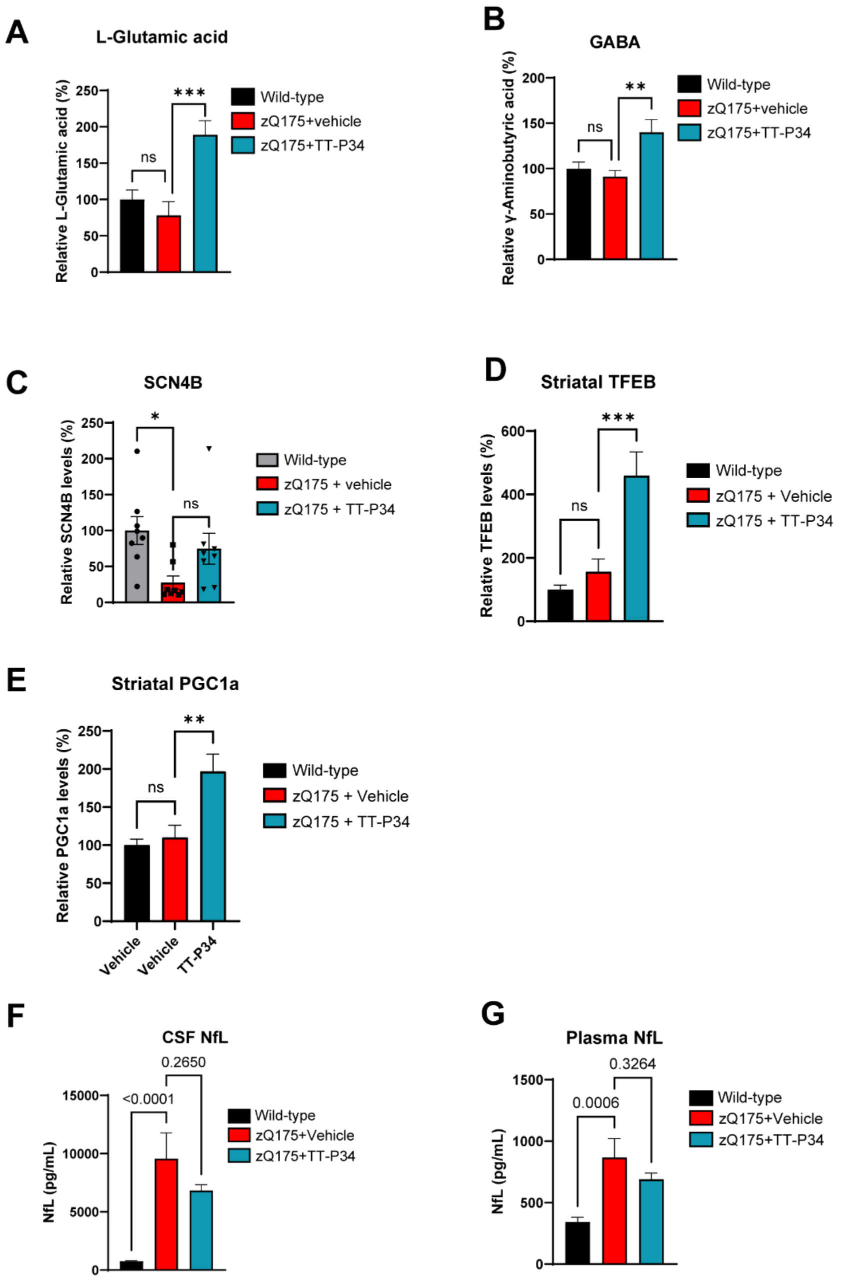


**Sup. Fig. 6: Post-mortem changes in metabolites and proteins of the striatum and biofluids from TT-P34 treated zQ175 mice**

**(A)** Relative L-Glutamic Acid and **(B)** GABA levels in the striatum of TT-P34 treated and vehicle-treated zQ175 versus wild-type measured by metabolomics (n=9). Relative SCN4B **(C)**, **(D)** TFEB and **(E)** PGC1a levels in the striatum of TT-P34 treated and vehicle-treated zQ175 versus wild-type measured by simple western (n=9). **(F)** Neurofilament Light chain (NfL) levels in cerebrospinal fluid and **(G)** plasma of zQ175 mice versus wild-type (n=18-20). Significance was calculated using ordinary one-way ANOVA. Mean ± SEM.

A

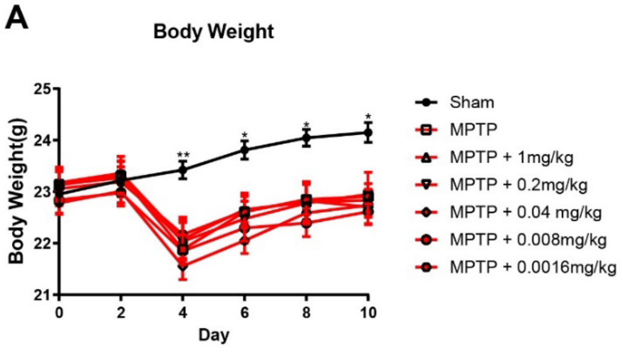


B

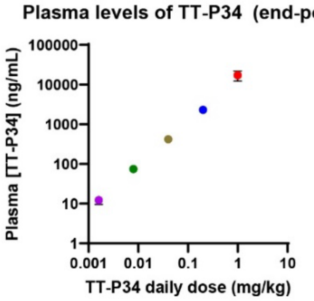


C

| Dose (mg/kg) | Plasma conc. (ng/mL) | Brain conc. (ng/g) (1%) | Brain free fraction (ng/g) (Fu brain: 0.0683) |
|---|---|---|---|
| 1 | 17143 | 171.43 | 12.00 |
| 0.2 | 2316 | 23.16 | 1.62 |
| 0.04 | 421 | 4.21 | 0.29 |
| 0.008 | 74.3 | 0.743 | 0.05 |
| 0.00016 | 12.3 | 0.123 | 0.01 |

**Sup. Fig. 7: Body weight and endpoint plasma concentrations of TT-P34 in MPTP mouse model**

**(A)** Bodyweight of MPTP-treated mice treated with once-daily subcutaneous administration of TT-P34 at different dosesand wild-type (WT) mice (sham = no MPTP treatment) throughout study (n=12). Significance was calculated using Nonparametric tests Kruskal-Wallis tests as data did not follow Gaussian distribution **(B)** Plasma endpoint levels of TT-P34 in MPTP-treated mice (n=12). **(C)** Plasma endpoint levels and calculated brain and brain free fraction levels of TT-P34 treated MPTP-treated mice. Mean ± SEM.

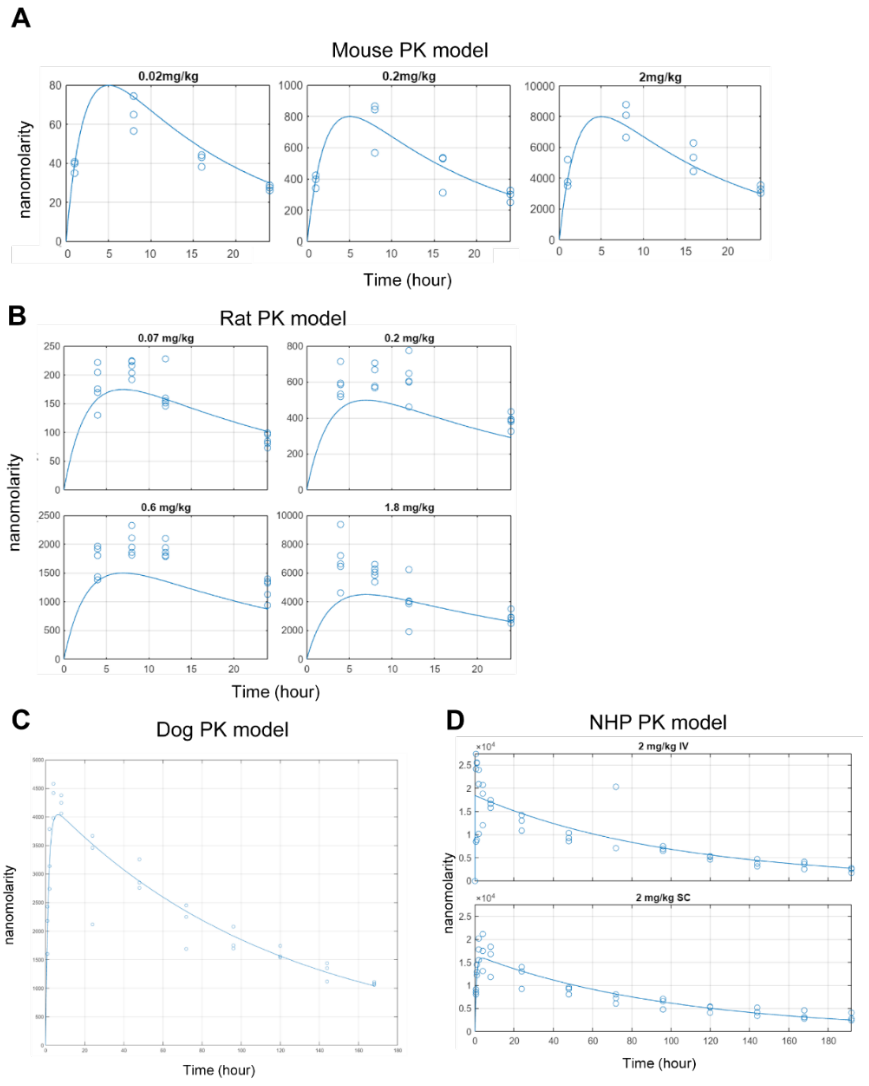


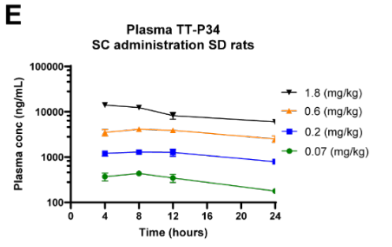


**F**

| Parameter | Estimate |
|---|---|
| Ksynth | 1856.6 |
| Kdeg | 19.76 |
| EC50 plasma free (nM) | 2 |

**Sup. Fig. 8: Pharmacokinetic models of TT-P34 plasma concentrations in mice, rats, dog and non-human primates**

**(A)** Pharmacokinetic model curve of TT-P34 in plasma at different timepoints and concentrations compared to observed concentrations (shown in dots) in mice (n=3) and in **(B)** rats following subcutaneous administration (n=3). **(C)** Pharmacokinetic model curve of TT-P34 in plasma at different timepoints compared to observed concentrations (shown in dots) in dogs following subcutaneous administration (n=3). **(D)** Pharmacokinetic model curve of TT-P34 in plasma at different timepoints compared to observed concentrations (shown in dots) in non-human primates following subcutaneous and intravenous administration (n=3).

### Supplementary Methods

#### Chloroalkane penetration assay

CAPA was performed as previously described [56-58]. The Hela cell line used for CAPA was generated by Chenoweth and co-workers to stably express HaloTag exclusively in the cytosol [59]. Cells were seeded in a 96-well plate the day before the experiment at a density of 4×104 cells/well. On the day of the experiment the media was aspirated, and Opti-MEM (100 µL) was added to the cells. CA-tagged peptide stocks in DMSO were diluted in Opti-MEM and serial dilutions of the peptides (1:1) were performed in a separate 96-well plate, ensuring the final DMSO concentration was kept consistent at 3%. Next, peptide solution (25 µL) was added to each well (various concentration range) and the plate was incubated for 4 h at 37 °C with 5% CO2. The contents of the wells were aspirated off, and wells were washed using fresh Opti-MEM (80 µL) for 15 min. The wash was aspirated off, and cells were chased using CA-TAMRA (5 µM, 50 µL) for 15 min, except for no-CA-TAMRA control wells, which were incubated with Opti-MEM alone (50 µL). The contents of the wells were aspirated and washed with fresh Opti-MEM (80 µL) for 30 min. After aspiration, cells were trypsinized (20 µM), resuspended in PBS containing 2% FBS (180 µM), and analyzed using a benchtop flow cytometer counting 2000 GFP+ events (cells) in up to 240 s per well. Obtained mean red fluorescence intensity data was normalized based on no-CA-tag wells (high fluorescence, 100% penetration) and no-CA-TAMRA wells (low red fluorescence, 0% penetration). A small molecule control CA-Trp-NH2 (CP50 = 38 ± 1 nM, n = 5) was included on each plate for referencing and quality control. Assay performance was controlled by Z' value determination resulting in Z' = 0.930 (n = 5). Viability of HeLa cells was measured as% GFP+ cells and plotted along the normalized mean fluorescence intensity.

#### Lysate stability assay

Hela cell lysate was freshly prepared by lysing the appropriate amount of cells (corresponding to CAPA conditions – lysate of 4×104 cells/replicate) in Tris buffer containing 1% triton-X at either pH 7.4 or 5.5. Assay matrix (90 µL) was preheated to 37 °C (5 min) and spiked with 10 µL (500 µM stock) of peptide of interested to generate a final concentration of 50 µM. Time point samples were taken at 0 h (immediately after spiking the lysate) and 4 h but quenching 40 µL of assay matrix with 40 µL of 6 M urea. After vortexing, the samples were incubated on ice for 30 min. Next, 40 µL of 20% TCA in acetone (w/v) were added and the samples were incubated for 2 h on ice. Before LCMS and UPLC analysis, samples were spin down at 13.400 rpm for 5 min. LCMS methods as outlined above were used to analyze the samples by injecting 2 µL. UPLC analysis was performed by using the methods outlined above by injecting 20 µL of sample.

**LC/MS detection of peptides in plasma, brain or CSF**

For plasma and CSF samples, 5µl of working solution was spiked in 45µl of plasma diluted 1:2 with HEPES or CSF diluted 1:2 with plasma. 50µl of sample were extracted using 300µl of MeOH containing Leu-Enk or Rolipram as Internal Standard. Centrifugation 4000rpm for 15min at 4°C. 100 µl of supernatant were transferred into 96 dwell plate containing 150 µl of Water and 20µl was injected into the UPLC system. For measurement in brain homogenates Solid-Phase Extraction (SPE) was used in order to clean and concentrate samples. 10 µl of working solution was spiked in 90 µl of brain homogenized 1:5 with MeOH. 50 µl of supernatant was diluted with 50µl of NH3 5% and loaded in Waters Oasis MAX SPE and wash with Ammonium Formiate 20mM and MeOH. For elution 50µl of TFA 1% ACN/Water 75/25 was used. 18µl of sample was injected into the UPLC system coupled with zenoTOF (High Resolution mass Spectrometer).

**Plasma & brain binding**

Compounds were dissolved in in water solution 0.01M $NH_4HCO_3$ pH 7.4 to give 20mM solution. Further dilutions (1:5) was prepared to obtain working solution, which was spiked into plasma obtaining a finale concentration of 20 µM. From initial spiking solution, compounds were immediately extracted and used to calculate recovery (T=0). An equilibrium time of 5 hours was otherwise used to assess plasma binding. 150 µL of PBS was dispensed in half-well 96 well dialyzer apparatus (HTDialysis LLC) and 150 µL of spiked plasma was loaded in other half-well. At end of equilibrium period (5 hours at 37 °C under shaking), 50 µL of dialysed plasma was added to 50 µL of corresponding PBS, and vice versa for buffer, to obtain same buffer to matrix ratio. The compounds were then extracted by protein precipitation with 500 µL of Methanol containing Rolipram (20 ng/mL) and Leu-Enk as internal standard (25 ng/mL) and centrifuged for 15 min. at 4000 rpm. Supernatants were collected and evaporated under nitrogen stream and resuspended in 100 µL of MilliQ water 25 % Methanol. Samples was injected into 6500 LC-MS/MS system. Protein binding was determined with following formulas: Afu=PBS/Matrix, where Afu = apparent fraction unbound,

PBS = analyte in buffer compartment, Matrix = analyte in plasma or brain compartment. The fraction unbound corrected (fucr) was calculated as fucr=(1/D)/((1/Afu-1)+1/D), where D is dilution factor (1 for plasma). The %Binding=(1-fucr)x100. Recovery rate was also calculated (not shown).

##### PK Model and Prediction of Human Parameters

Pharmacokinetic (PK) parameters were initially estimated independently for four animal species: mouse, rat, dog, and non-human primate (NHP). Due to the absence of intravenous (IV) data for mice and dogs, their bioavailability (F) was set to 1 and hence all parameter estimates are /F. For rats and NHPs, where both subcutaneous (SC) and IV data were available, bioavailability was quantified alongside clearance (CL), volume of distribution (Vd), and the first-order absorption rate constant (ka) through PK modelling.

Model development and parameter estimation was carried out using the SimBiology and Statistics Toolbox in Matlab 2023b. The established one-compartment model was selected based on its ability to effectively describe the pharmacokinetic data. A graphical representation of this model, illustrating drug administration via IV, oral (PO), or SC administration routes is provided in Figure 1.

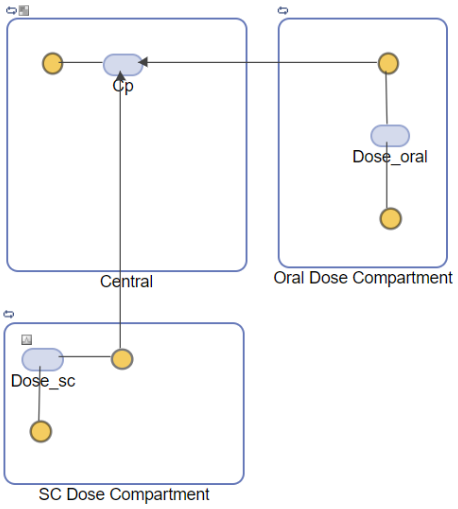

**Figure 1: One-compartment PK model. Drug delivered via IV (directly into Central compartment), PO or SC**

**doses.**

The parameters were directly parameterised per kilogram of body weight, thus simplifying the model application across various species without necessitating additional equations to adjust for body weight.

Following parameter estimation, allometric scaling was employed to extrapolate the animal PK data to predict human pharmacokinetic parameters, assuming animal weights as Table 1. This approach successfully predicted human PK parameters from mouse, rat, and NHP data for Semaglutide (REF), providing a valid foundation for predicting human clearance (CL) and volume of distribution (Vd) for TTP34.

**Table 1: Animal Weights Used for Allometric Scaling to Predict Human Pharmacokinetic Parameters.**

| Species | Weight (kg) |
|---|---|
| Mouse | 0.02 |
| Rat | 0.25 |
| Non-human Primate | 10 |
| Human | 70 |

#### Human Dose Prediction

Using the human PK parameters estimated from the allometric scaling, the dose of TTP34 for human use was predicted under two different assumptions regarding drug concentration targets:

1. Targeting the pharmacokinetic (PK) trough concentration at the level associated with maximum disease effect.
2. Targeting the average steady-state concentration (Cave) at the level associated with maximum disease effect.

These assumptions were based on the maximum pharmacodynamic (PD) effect observed at a dosage of 1 mg/kg in mouse models. The Cave for this dosage in mice, calculated using the mouse PK parameters, was determined to be 4850nM. With a mouse unbound fraction in plasma (fu) of 1.45%, this concentration corresponds to a free plasma concentration of 70nM. The mouse brain partition coefficient ($K_{puu}$) was 0.05, resulting in an effective free drug concentration in brain tissue of 4nM.

To translate these effects to humans, we assumed a 1:1 translation from mouse PKPD. A human dose was calculated to achieve similar brain exposure, based on the $K_{puu}$ values obtained from mouse, rat, and NHP studies. The $K_{puu}$ values for mouse (0.05), rat (0.5) and NHP (1) were considered. The NHP model, being physiologically closest to humans, was used to estimate a human dose range.

#### Application and Refinement of PBPK Model

We utilised a previously published PBPK model originally developed for Acetaminophen, which accurately predicted the PK in plasma and cerebrospinal fluid (CSF) both in rats and humans. We first replicated the existing model with Acetaminophen to confirm its accuracy by reproducing the PK profiles in plasma and CSF. Successful replication provided the necessary validation to proceed with adaptations for TTP34.

The model was refined to incorporate the PK parameters of TTP34 obtained from rat studies. Furthermore, the clearance parameter from plasma to brain ECF was estimated using total brain PK concentration data from rats. This integration enabled adaptation of the model accurately toTTP34, reflecting its unique PK properties. Figure 2 illustrates the PBPK model refined for TTP34.

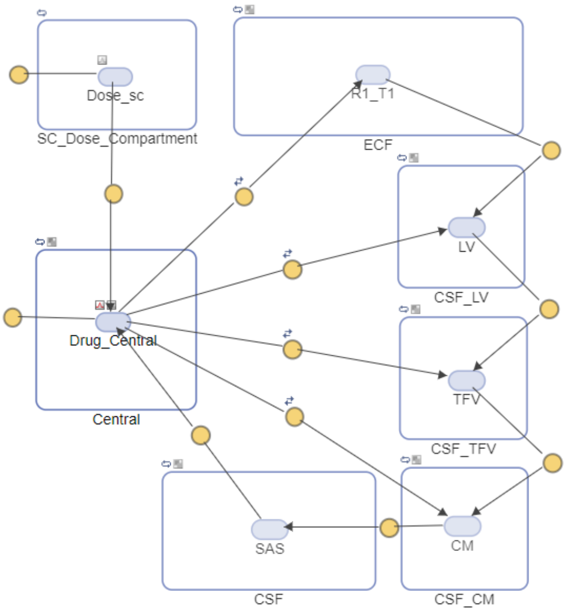


**Figure 2: Graphical representation of the PBPK model adapted from [70] for TTP34 to describe the intra brain distribution in rat and human.**

Using the refined model, simulations were performed at four different dose levels in rat, demonstrating good concordance with observed data for chronic plasma, CSF, and total brain concentrations. Further simulations were conducted using the NHP PK parameters, with a weighted average of rat and human parameters derived from the published model. This approach maintained the model's remarkable accuracy in predicting the PK profiles across species.

Finally, the human doses previously predicted—those necessary to achieve targeted trough and targeted $C_{ave}$—were simulated using the refined PBPK model using the extrapolated human PK parameters and the model parameters obtained from the published model [70]. This step aimed to predict human CSF subarachnoid space (SAS), total CSF, and free brain concentrations of TTP34, providing critical insights into the expected human pharmacological profiles, aiding in the estimation of efficacious and safe dosing regimens.

**Software Utilization**

The development and analysis of the PKPD were conducted using SimBiology, a module within Matlab specifically designed for building and simulating biological systems, including drug kinetics. This environment provided tools for iterative model fitting, parameter estimation, and model simulations, facilitating rigorous validation of the model parameters against the experimental data.
